# Spatially defined axonal guidance in neural organoids with micropatterned microfluidic channels

**DOI:** 10.64898/2026.04.30.721979

**Authors:** Ariana Cisneros, Maryam Moarefian, Jens Duru, Katherine Karinicolas, Talia Goodman, Zaira Gonzalez, Anton Zatserklyaniy, Sawyer McKenna, Asia Anderson, Noah Wiliams, Gregory Kaurala, Estefania Sanchez, Ali Shariati, Mircea Teodorescu, Tal Sharf

## Abstract

Three-dimensional stem cell-derived neural organoids provide a promising platform for investigating early brain development and interregional circuit formation. Although co-culture of region-specific organoids into assembloids has enabled the study of cortical and subcortical interactions, these models lack directional specificity and spatial control, limiting their ability to recapitulate canonical circuit architecture. Here, we present a microfluidic platform for constructing directional and tunable interregional circuits while preserving anatomical distinction. This system, which we term “directoids” incorporates micropatterned polydimethylsiloxane (PDMS) microstructures to control uni- and bidirectional axonal growth between cortical and thalamic organoids. We observed a 70.4% success rate of axons traversing the full channel length in the permissive direction and reaching the opposing organoid, whereas no neurites successfully crossed the probative direction. These results demonstrate robust directionally bias in axon outgrowth and establish a scalable, reproducible strategy for controlling asymmetric connectivity between anatomically distinct neural organoids. Using high-density CMOS microelectrode arrays, we further validated directional tuning of extracellular action potential propagation within directoid microchannels, a feature not observed in straight-channel connectoid controls. Directoids also exhibited significant asymmetry in firing rates between channel entry and exit sites, consistent with engineered bias in signal flow. This provides an experimental paradigm for dissecting how anatomical connectivity and functional activity converge to shape neuronal networks. Together, these findings establish a microfluidic platform for investigating the mechanisms underlying hierarchical circuit formation, regional specification, and functional integration in developing human neural organoid models at cellular resolution not possible *in vivo*.

## Introduction

Through morphogenesis and self-organization^1–3^, the mammalian brain establishes large-scale circuit wiring and a functional connectome with conserved features across brain regions and phylogeny^4^. This connectome is marked by highly connected hubs and modular architecture^5^, formed through directionally weighted connections that span anatomically distinct brain regions^6^. Despite these conserved features, it remains unclear how developing brain regions interact during early stages of brain development, and how they acquire their regional specificity^7^. These early developmental events ultimately give rise to efficient and economically organized computing regimes in the brain^8,9^. Large-scale single cell transcriptomics has revealed how gene programs influence cell fate decisions in the developing human brain^10^, but extrinsic mechanisms influencing regional brain formation, especially within the cortex, are not fully understood^11^.

Over the last several decades two prominent theories have emerged to explain cortical development and specialization. The ‘*protomap hypothesis’* posits that genetic programs intrinsically define regional identity^12,13^, while the ‘*protocortex hypothesis’* states that early cortical neurons are generalists where terminal identity is shaped by activity dependent patterning from subcortical afferents, for example from the thalamus^7,14,15^ (Fig. 1A,B). Testing these hypotheses *in vivo* has proven difficult, as developing brain regions cannot be isolated without disrupting their cellular context^7^. Furthermore, sensory input relayed by thalamic afferents^16^, which shape terminal cortical identity additionally, further confounds intrinsic versus extrinsic contributions to early circuit dynamics^17^.

**Figure 1.**
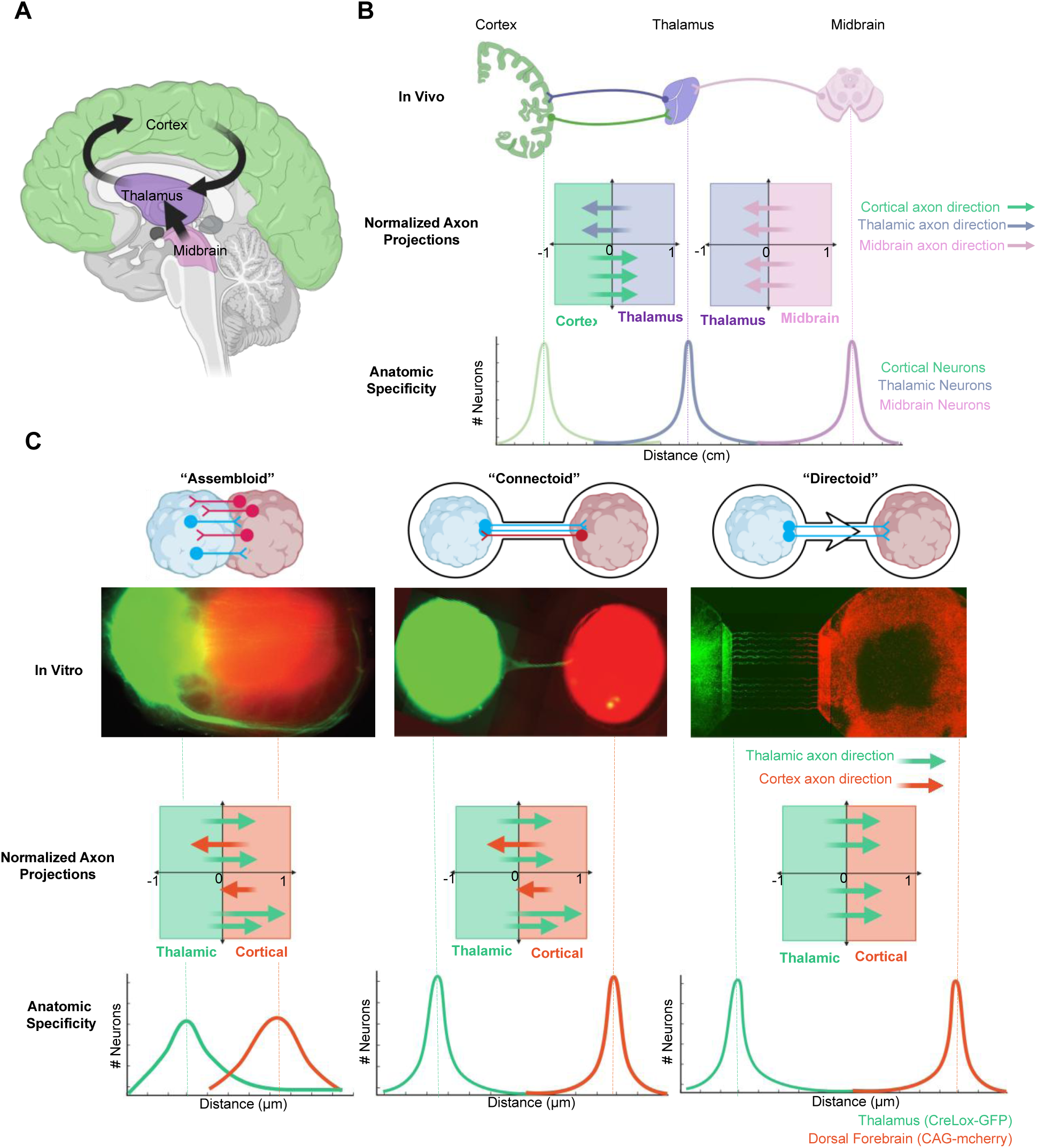
Bioengineering directional circuits across cortical and subcortical tissue *in vitro*. **(A)** Schematic diagram illustrates that both anatomical organization and directional routing of axon tracts are key features brain connectivity (e.g. the corticothalamic loop). **(B)** *In vivo* axon projections maintain high regional specificity with directionally enriched neurite trajectories between cortex, thalamus, and other subcortical structures (e.g. midbrain). **(C)** Top, Comparison of *in vitro* models: conventional neural assembloids exhibit random, overlapping neurite outgrowth and cellular migration that leads to ambiguous anatomical organization. Connectoids (straight microchannel geometry) maintain anatomic specificity but lack directional guidance for neurite outgrowth; directoid (asymmetric microchannel geometry) microstructures maintain regional anatomic specificity while directionally guiding neurite outgrowth between organoid regions. Middle, representative fluorescent images of human cortical (CAG-mCherry, red) and thalamic organoids (creLox-GFP, green) grown from cell lines engineered to express fluorescent proteins for an assembloid (left, 46 days *in vitro*), connectoid (middle, 51 days *in vitro*) and directoid (right, 55 days *in vitro*). Bottom, diagram overview of axon projections directionality and anatomic specificity curves indicating directional bias.

Recent advances in single-cell multi-region analyses have enabled comparative molecular profiling of homologous cell types across early human brain regions^18^, supporting a revised model of cortical development: broad proto-regions emerge early and are later refined into functionally specialized areas via activity-dependent mechanisms. However, these insights are drawn from static postmortem tissue, limiting our ability to study dynamic processes like axon guidance, synapse formation, and circuit integration. A fundamentally new approach is needed to understand how specialized microcircuits emerge and mature through activity-dependent changes in cell identity, within the broader cytoarchitecture that ultimately gives rise to the reciprocal wiring of the corticothalamic loop^19^.

The rise of three-dimensional, stem cell–derived brain models, known as organoids, has enabled the *in vitro* assembly of developing brain structures that recapitulate key aspects of anatomical organization and cellular diversity^20–26^ as well as the functional network dynamics observed in the developing human brain^27–31^. More recently, assembloids, formed by fusing region-specific organoids, have been used to model interactions between cortical and subcortical regions^32–36^, and the genetics of corticothalamic wiring^37^. Despite these advances, current assembloid techniques rely on passive fusion of tissues, often resulting in ambiguity regarding regional identity due to post-fusion proliferation and cell migration^38,39^ (Fig. 1C). To overcome these challenges, emerging bioengineering approaches aim to guide tissue structure and function with greater control and precision^40^. These include acoustic manipulation^41^, bioprinting^42^, and integration with microfluidic systems to promote functional maturation and establishment of local and long-range axonal connections^43–45^.

Here, we present a novel bioengineering approach that uses asymmetric microchannels to guide axonal polarity between anatomically distinct, brain-region-specific organoid tissues. Our method builds on work by Taylor *et al*^46^, Peyrin *et al*.^47^, Renault *et al*.^48^, Holloway *et al*.^49^, and Girardin *et al*.^50^, which leverage the finite bending stiffness of neurite growth cones and edge guidance along microchannels^51^ to route unidirectional neurite outgrowth in two dimensional neural cultures^52^.

By physically constraining organoids in custom microfluidic compartments, we developed a tunable platform that directs neurite outgrowth from subcortical to cortical organoids through intrinsic axon guidance programs and without relying on complex molecular guidance cues^53,54^. Our optimized geometric design eliminated axon traversal in non-target directions by reducing traversal speed fourfold while achieving a 70% success rate for axons in target directions. To confirm unidirectional axon propagation between our directional organoids we used high density complementary metal-oxide-semiconductor (CMOS)-based microelectrode arrays (MEAs)^55^ to recorded spontaneous action potential propagation across axons in both asymmetric and straight microchannels. We observed unidirectional propagation in asymmetric microchannels, while straight microchannels displayed bidirectional action potential propagation. Finally, implementation of our device led to a significant increase in the relative spike rate frequencies at the entrance relative to channel exits for directional microstructures (54%) compared to conventional straight microchannel (3%) configurations. This design creates a competitive advantage for neurites following permissive paths, establishing a new experimental paradigm to investigate how subcortical inputs shape early cortical identity and function prior to the onset of sensory experience.

## Results

### Directional steering of axonal tract extension from a neural organoid

To create tunable directional axon tracts between neural organoids, we developed geometrically defined microchannels which exploits neurite edge guidance and asymmetric microchannel geometries to direct, trap, or redirect axonal growth polarity^46–49^. Our microstructure utilizes alternating channel traps to inhibit growth in prohibitive directions while allowing growth in permissive directions (Fig. 2A). Directoid microstructures were fabricated by casting poly(dimethylsiloxane) (PDMS)^56–58^ on a custom photolithography mastermold fabricated using conventional photolithography techniques (Fig. 1A, Supplementary Fig. 1).Channel heights of <3.5 μm effectively filtered for growing neurites while excluding neural progenitor somas (Fig. 2B). Channel heights were validated using a surface scanning profilometer and fluidic viability assessed by flowing fluorescent beads through microchannels under vacuum (Supplementary Fig. 1B).

**Figure 2.**
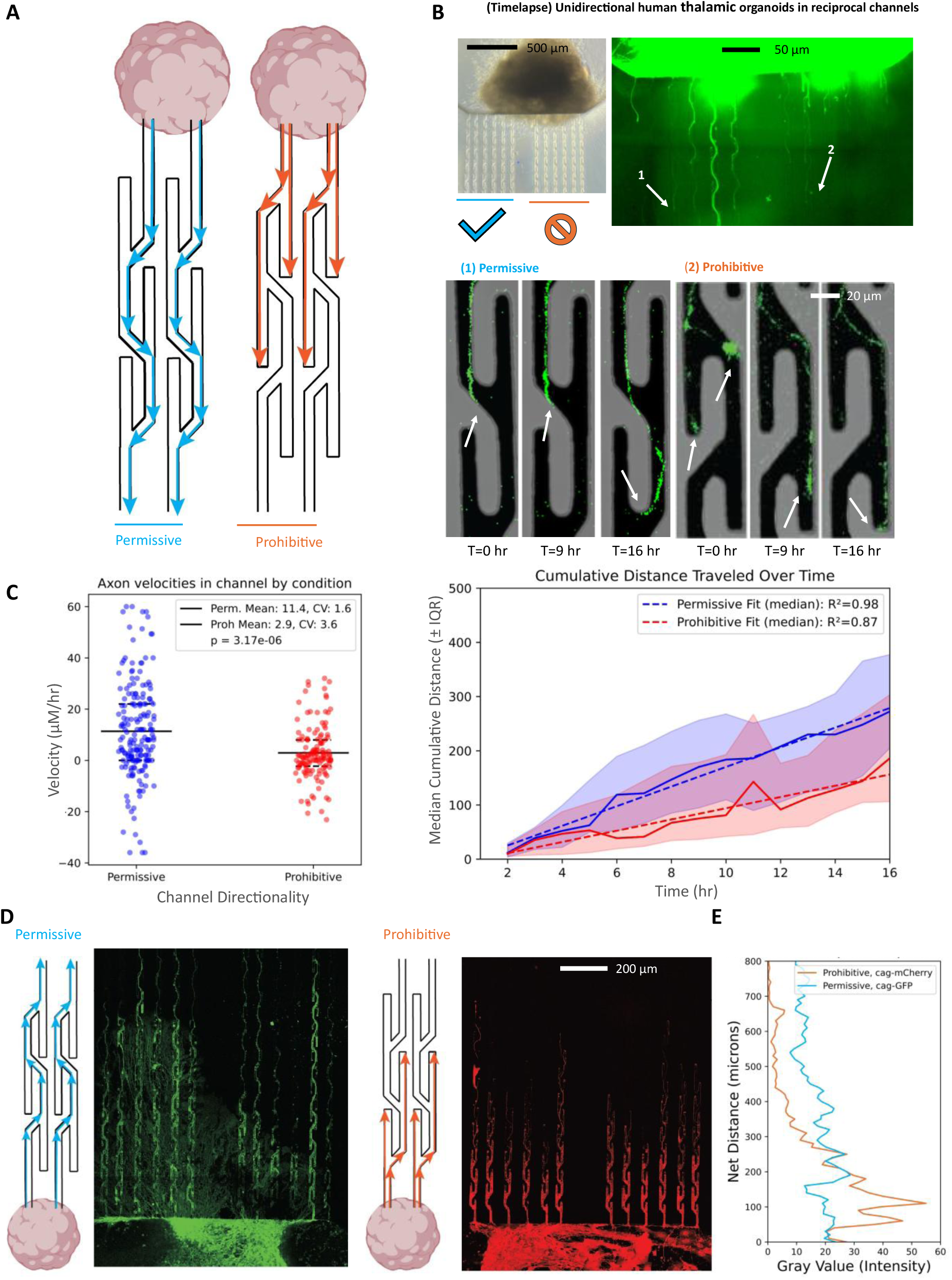
Engineered microenvironments guide directional axon outgrowth from human thalamic organoids. (**A**) Schematic of a bidirectional microchannel platform for engineering directional axon circuits. Asymmetric geometries create either permissive (blue) or prohibitive (orange) environments to bias axon trajectory. (**B**) Time-lapse imaging of GFP-labeled neurites from human thalamic organoids demonstrates robust forward outgrowth in permissive channels, with rerouting or stalling observed in prohibitive channels. White arrows indicate axon extension over time. (**C**) Left, quantification of neurite dynamics from hourly imaging over 18 hours (*n* = 97 permissive tracks, n = 90 prohibitive tracks). Neurites in permissive channels extended at an average velocity of 11.4 µm/hr, compared to 2.9 µm/hr in prohibitive channels (Student’s t-test, *P* = 3.17 × 10⁻⁶; CV: permissive = 1.6, prohibitive = 3.6). Right, median distance traveled is plotted ± interquartile range (IQR), with linear regression fit over time (*R*²: permissive = 0.98, prohibitive = 0.87). (**D**) Endpoint projections after 15 days in microchannels reveal continuous outgrowth in permissive environments and confinement in prohibitive geometries. Right panel: Plot of net neurite distance as a function of normalized fluorescence intensity along the 900 µm long migration axis. Permissive distributions (green) exhibit broader neurite spread, while prohibitive conditions (red) show sharp attenuation and clustering near the channel entrance. (**E**) Quantification of overall pixel intensity for unidirectional permissive trajectories (creLox-GFP) demonstrates near millimeter extension, whereas unidirectional prohibitive trajectories (CAG-mCherry) demonstrate diminished growth over distance traveled.

Our initial design was inspired by the concept of the “axonal diode”^46–49^. We found that organoids, which contain a higher density of neurons in a three-dimensional volume compared to two-dimensional cultures with similar cross-sectional area, can still extend neurite processes through quasi-two-dimensional microchannel interfaces with multiple triangular sharped axonal-diode geometries as shown in previous work (Supplementary Fig. 1A). Optimization for seeding human organoids into microwells required shorter channels to prevent early neural progenitor cell migration into microchannels, which was not observed in mouse cultures (Supplementary Fig. 3E).

Initial microstructure prototyping utilized mouse embryonic stem cell (ESC) derived cortical organoids, adapted from Eiraku *et al*.^20^. Mouse brain development is accelerated by approximately 7-fold relative to the human brain^7^ allowing for rapid prototyping to take place over weeks rather than months. Here, we demonstrate that a pool of immature neuronal progenitors is required to promote axon extension into microchannels (7 days *in vitro*), whereas when neurites are already formed within mature organoids (>21 days *in vitro*), their processes will not extend into the channels (Supplemental Fig. 2D). This effect is mirrored in human organoids from 30 days *in vitro* to 45 days *in vitro* (Supplemental Fig. 2E).

We performed longitudinal live-cell imaging to capture growth cone dynamics over a 16-hour period (Fig. 2B, top) to quantify axon traversal through microchannels. We generated stem cell derived thalamic organoids^26^ using a protocol developed by Xiang *et al*.^33^ from an H9 cell line expressing green fluorescent protein (GFP) under the control of the synthetic CAG promoter^59^. Growth cones preferentially follow the straightest trajectory available^46–49,60^, which informed our alternating straight trap design. Our imaging revealed that when growth cones traversed the microchannels with upward branches, defined as the *prohibitive* orientation, they are routed into dead end traps and are forced to reroute to continue growth (Fig. 2A, left). In contrast, when growth cones traverse the microchannels with downward branches, or the *permissive* orientation (Fig. 2A, right), the axon easily bypasses the traps.

We next quantified distance dependent neurite outgrowth within our microstructures. Live-cell imaging of our fluorescently labelled organoids revealed complex spatiotemporal dynamics of axon growth within the confines of our microstructures, highlighting axonal branching and active exploration of growth cones in geometrically defined environments (Fig. 2B, bottom). Live imaging was utilized for velocity analysis and calculated by manually tracing hourly neurite extension across three different experimental microstructures. One microstructure contained a single thalamic organoid with bidirectional channels, and two bidirectional microstructures contained a dorsal forebrain CAG-mCherry tagged organoid and a thalamic creLox-GFP tagged organoid (*n* = 5 organoid biological replicates). Single axon growth cones were tracked manually using NeuronJ’s axon tracing tool in FIJI to follow individual neurite tracks over a 16-hour live imaging session. We calculated velocity as the mean neurite displacement as a function of time (Fig. 2C, right). Neurites in permissive trajectories grew at an average rate of 11.4 μm/hr±0.18 μm/hr, compared to just 2.9 μm/hr ±0.10μm/hr in prohibitive trajectories, a fourfold decrease in velocity (Fig. 2C, left, student’s t-test: *P =* 3.16×10^−6^*, n* = 97 neurites tracked across 5 organoids). Median rate of outgrowth scaled linearly as revealed by linear regression (Fig 2C, right. R²: permissive = 0.98, prohibitive = 0.87). The rate of axon extension is consistent with prior *in vitro* studies of neurite growth extension in human iPSC *in vitro* cultures^61^. To examine how our microchannel geometry impacted long-term neurite outgrowth distributions within our directional microstructures, we tracked neurite outgrowth from human cortical and thalamic organoids daily on a fluorescence microscope. We observed consistent neurite outgrowth into the channels within approximately 4 to 5 days after seeding organoids that were 30 days *in vitro*.

The distribution of neurite lengths was measured for eight organoid replicates in directoid microstructures with directional microchannels and four organoid replicates in control microstructures with straight channels. Over the course of longer time durations, we observe clear distinction between axon length as a function of distance across the 900 µm traversal distance for the permissive and prohibitive microchannel designs, with axons completely stalling within a few hundred microns in the prohibitive direction. Meanwhile axons in the permissive direction spanned close to millimeter scale distances (Fig. 2D,E), demonstrating our approach allows us to control and precisely measure long-range directional axonal outgrowth.

### Cellular characterization of cortical and thalamic organoid regional identities

To confirm the identities of our cortical and thalamic organoids within microfluidic device we performed immunohistochemistry to validate the presence of cell types across key developmental time points. Visualization of individual organoid processes was performed by generating dorsal forebrain and thalamic organoids using two distinct ESC lines expressing GFP and mCherry, respectively (see Methods). Our cortical organoids were generated following the methods developed by Yoon *et al*.^62^ to generate tissue of prominent forebrain identity. For dorsal forebrain organoids we confirmed that our early developing cortical organoids had a robust pool of SOX2 positive neuronal progenitors at 33 days *in vitro* (Fig. 4A, top left). Given the presence of early progenitor pool, we next sought to confirm the presence of terminally differentiated neurons. We confirmed both the presence of MAP2 positive neurons well as specific brain region cortical patterning demonstrated expression of the dorsal forebrain marker FOXG1 at 43 *days in vitro*, where we observed the formation of circular rosette-like structures composed of the proliferating stem cells and early neural progenitors that recapitulate neural tube formation^26^, (Fig. 4A, bottom left). Next, we validated the patterning and viability of our thalamic organoids which we generated following the methods of Xiang *et al*^33^. (see Methods). We confirmed development of PAX6-positive neural progenitors in our thalamic organoids at 33 *days in vitro* (Fig. 4B, top right). At 48 *days in vitro*, we visualized broadly expressed thalamic projection marker FOXP2, and the pan neuronal marker MAP2 to confirm presence of a maturing neuronal population. (Fig. 4B, bottom right). Together, these results confirm the presence of key cell types required to establish an organoid-based model of the corticothalamic interregional connectivity.

**Figure 3.**
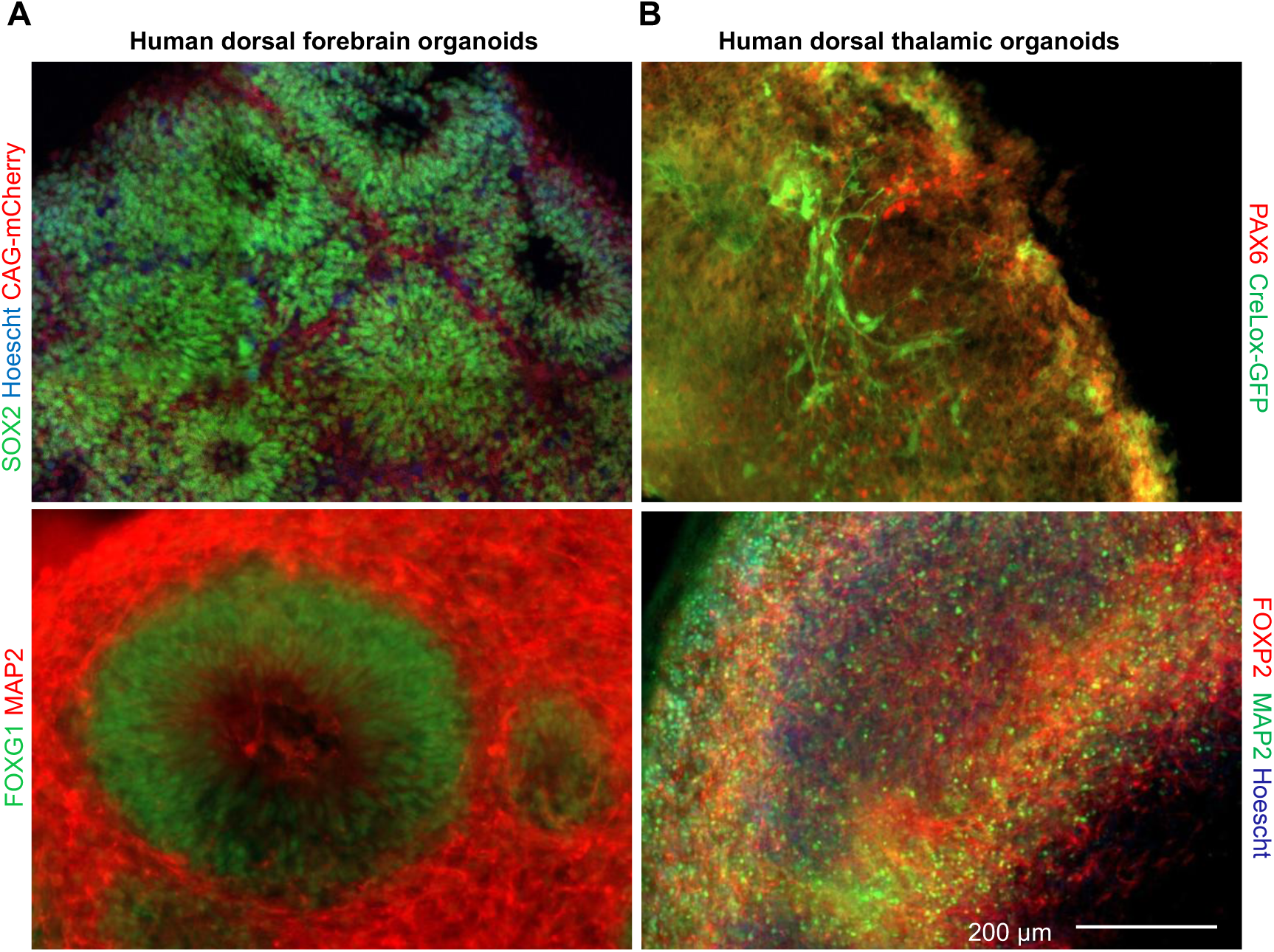
Characterization of cell types and regional identity of human cortical and thalamic organoids. (**A**) Representative immunostained images of human cortical organoids of dorsal forebrain identity. A subset of cells expresses the stem cell marker SOX2 (top left) at 33 days *in vitro*. The dorsal forebrain marker FOXG1 and somadendritic compartment marker MAP2 at 43 days *in vitro* (bottom left), revealing the expansion of deep-layer cortical neurons. Cortical organoids in fluorescence experiments express red mCherry under the CAG promoter **(B)** Representative immunostained images of thalamic organoids reveal the presence of a PAX6 positive neuronal progenitor pool at 33 days *in vitro* (red, top left). FOXP2 is present at DIV48 as a broad thalamic marker as well as MAP2 positive neural marker (bottom right). Thalamic organoids in fluorescence experiments express GFP under the creLox promoter, scale bar 200 μm.

**Figure 4.**
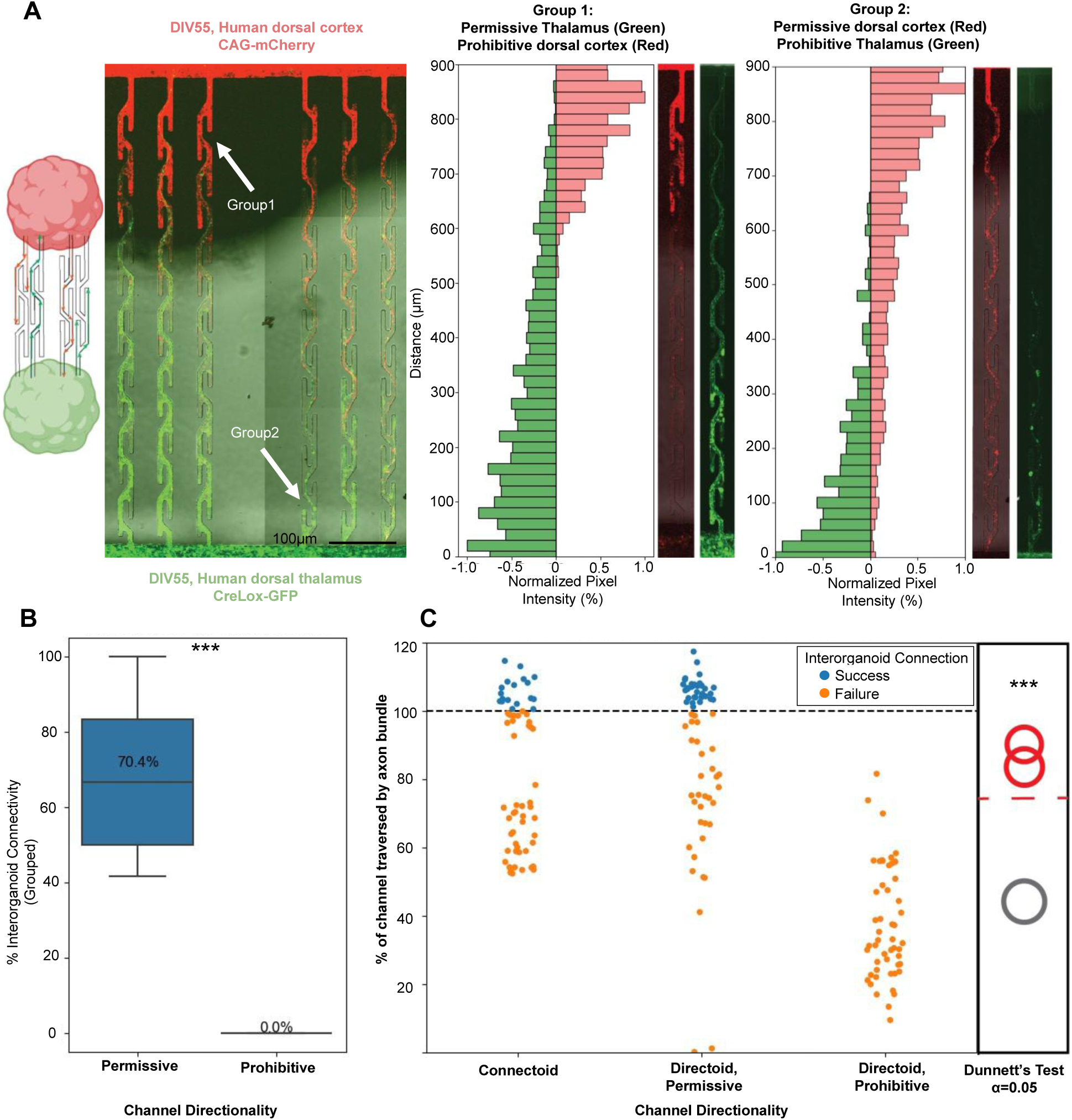
Engineering directional connectivity between human cortical and thalamic organoids. (**A**) Left, schematic diagram of a directoid device to establish reciprocal connectivity between cortical and thalamic organoid neurite projections. Right, representative image of axons outgrowth from human cortical (red) and thalamic (green) organoids within asymmetric microchannels at 55 days *in vitro* expressing mCherry and GFP respectively. Channels on the left-hand side (group 1) are prohibitive to cortical (red) to neurite growth from the top down and permissive thalamic organoid (green) neurite growth from the bottom up. Channels on the right side (group 2) have channel orientation reversed and prohibit neurite growth from thalamic organoids (green, bottom) and permit cortical neurite growth (red, top). Right, individual microfluidic channels are shown with neurites extending along their length. These examples refer to the channel locations identified by two arrows in the left panel. Histogram plots show the summed fluorescence intensity for each red and green colors in the images. Neurites in the permissive direction span the length of the channels for both the cortex (top-to-bottom) and the thalamus (bottom-to-top). **(B)** Quantification of organoid innervation success across 60 channel conditions. In permissive configurations, ∼70.4% of paired microenvironments support successful inter-organoid axon crossing, while prohibitive channels result in 0% span of the full channel length (*P*<0.0001, Mann Whitney U test) **(C)** Percent of each microchannel traversed by neurites in permissive, or prohibitive geometries. Control data refers to straight channels where organoids we placed on one side to measure outgrowth as a function of distance without curvature or traps. Axons that traversed the full channel length are shown in blue, while failures (orange) stall at intermediate positions. Summary analysis (right panel) shows significant differences in the distribution of neurite traversal lengths in permissive versus prohibitive orientations relative to the reference organoid using Dunnett’s test (*P* < 0.0001 One Way ANOVA, α = 0.05, *n* = 30 microstructure channels per condition).

### Directing axonal projections between cortical and thalamic organoids

Having established directional control of neurite outgrowth in individual organoids within our microfluidic channels and generated both cortical and thalamic organoids, we next combined these components into a single platform. To do this, we designed a microfluidic chip containing reciprocally oriented directional channels to direct axon projections between cortical and thalamic organoids (Fig. 4A, Supplementary Fig. 1B). To quantify how the directional microchannels routed neurites, we used cortical organoids expressing red fluorescent protein (mCherry) and thalamic organoids expressing green fluorescent protein (GFP). These organoids were positioned at opposite ends of the microfluidic device. Cortical and thalamic organoids were seeded into device wells at 30-35 days post aggregation, referred to as days *in vitro*. We monitored neurite outgrowth with a fluorescent microscope, imaging axons from the cortical (red) and thalamic (green) organoids (Fig. 4A), and quantified the distance traversed along each microchannel. Neurite outgrowth was tracked over several weeks, and tissue was fixed at 55 days *in vitro* for high-resolution confocal imaging (see Methods).

Analysis revealed that neurites from both cortical (Fig. 4A, left top) and thalamic (Fig. 4A, left bottom) organoids preferentially follow the permissive orientation of the directional microchannels. For example, red axons from the cortical organoid placed at the top of the device were unable to traverse down the left-side channels, where growth cones encountered dead ends. In contrast, neurites on the right-side permissive channels readily extended across the full length. A mirrored pattern was observed for green neurites from the thalamic organoid placed at the bottom of the device. To quantify these observations, we measured cumulative fluorescence intensity across the red and green channels to assess neurite outgrowth within individual microchannels (Fig. 4A, right). For microchannels 900 μm in length, we observed a 70.4% success rate of axons traversing the full channel length in the permissive direction and reaching the opposing organoid. In contrast, no neurites (0%) successfully traversed the full-length of the prohibitive direction within any of the directoid devices (Fig. 4B).

To assess reproducibility, we repeated the live imaging experiment across 5 microfluidic directoid devices containing a total of 30 microchannels. We observed a significantly greater axonal traversal in the permissive direction relative to the prohibitive direction (Fig. 4C; *P* < 0.001, Dunnett’s test, α = 0.05). As a control, we evaluated neurite outgrowth between single organoids placed at either end of straight microchannels to determine whether growth success depended on distance. We found no significant difference in outgrowth between straight control channels and the permissive directional channels (Supplementary Fig. 3). Based on this experimental data we conclude that the asymmetric geometry of the prohibitive channels, not the path length, limits long-range neurite extension in our directoids.

### Directional microchannels directionally bias action potential propagation between organoids

To assess directional action potential propagation in microstructures between two organoids, we attached our microstructures to the planar recording surface of high-density CMOS-based microelectrode arrays^55^ (see Methods). To confirm action potential propagation within microchannels between two organoids, we initially routed recording electrodes within the microchannels, used impedance scanning to assess PDMS adhesion to the planar surface (Supplementary Fig. 4) using a custom script from Duru *et al.*^63^. Output files were used to create custom routing configurations for each microstructure. Here, we used mouse embryonic stem cell (ESC)-derived thalamic organoids to characterize the directional properties of action potential propagation in our microchannel designs. We tested 4 connectoid microstructures (straight microchannels) and 4 reciprocally connected directoid microstructures (directional microchannels) on the microelectrode arrays. We next utilized the transmission line properties of action potential propagation to map co-occurring spike events with high fidelity and short latencies^64,65^ across the length of our microchannels (see Methods). Mapping co-occurring spike events revealed consistent unidirectional action potential propagation at velocities of approximately 0.65 mm/ms (Fig. 5A,B), consistent with previous reports of axonal conduction velocities in stem cell-derived neurons^60,64–66^. We observed consistent action potential propagation velocities in both connectoid and directoid devices. However, in directoid structures, propagation events were detected only in the permissive growth direction and not in the prohibitive direction (Fig. 5C, top right). In contrast, propagation events in straight channel connectoid devices were detected in both directions and equal spiking frequency was observed on both sides of the microstructure (Fig. 5C, top left). However, directoids showed a significant increase in spiking frequency near the exit side of the permissive channels. This difference allowed us to characterize directionality as the ratio between the firing rate observed in the exit quartile versus the entry quartile, which we refer to as the flux ratio. Straight microchannels exhibited a flux ratio of 1:1, with symmetric firing rates at the entry and exit of the channels, whereas asymmetric channels had flux ratios > 2:1 comparatively. To further illustrate this feature, the mean firing rate in the last quarter of a directional microchannel was 7.7 Hz, compared with 2.6 Hz in the first quarter, yielding an approximately threefold difference (Fig. 5C, right). In contrast, a representative straight microchannel exhibited a mean firing rate of 2.1 Hz in the first quartile (Q1) and 2.2 Hz in the last quartile (Q4), yielding a flux ratio very close to one. We attribute this increase to an accumulation of neurites entering from the non-permissive direction, thereby increasing the number of detected action potentials near permissive channel exits relative to their entrances. Consistent with this interpretation, our microscopy assays confirmed that directional channel entries contained axonal processes only from the proximal organoid, whereas channel exits contained axonal processes from both the proximal and distal organoids (Fig. 4A).

**Figure 5.**
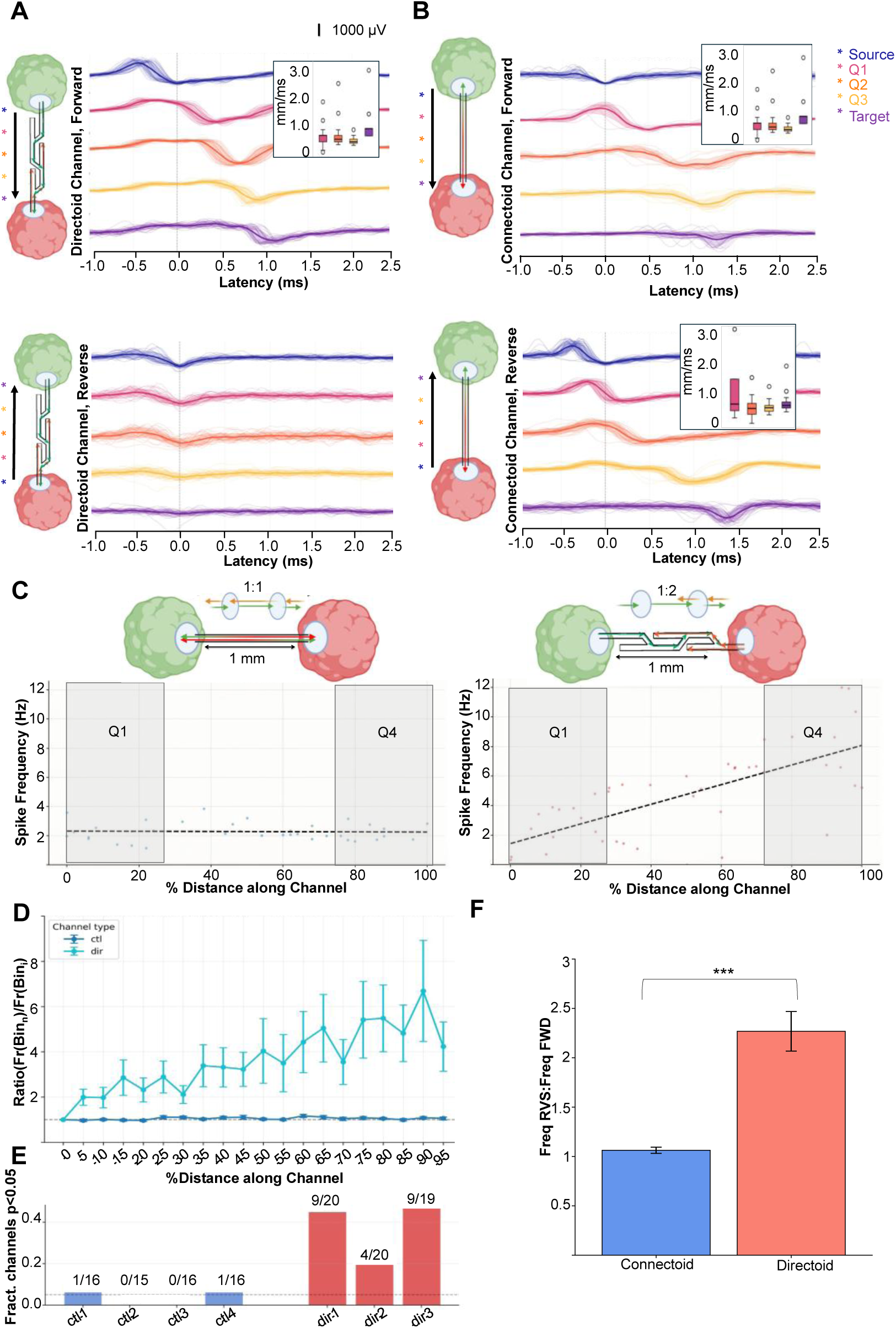
Directional control of action potential propagation between organoids. **(A)** Forward propagation of action potential spikes is present in permissive direction of asymmetric channels (Top Left). In the prohibitive direction (bottom left), no propagation signal was observed. **(B)** Action potential propagations are present in both directions for straight channel connectoid devices. Action potential velocity boxplots of firing events are shown with 0.65 mm/ms on average for all sampled positions where paired propagation events were observed. **(C)** Ratio of firing rate (FR) of last quartile (Q4) and first quartile (Q1) distance along a microchannel. Straight channels show spike frequency ratios of 1 across the channel, whereas directional microchannels spike frequency ratios increase across by 2.9 **(D)** Cross comparison of straight channel control microstructures vs directional channel microstructures. The FR-ratio increases across channel distance in directional microchannels, whereas FR-ratio remains flat across straight microchannels. **(E)** Comparison across four straight channel control microstructures and three directional microstructures reveal clear differences in firing-rate asymmetry when comparing channel entry and exit quartiles. 54% of directional channels show significant increases in firing rate (*n* = 3 organoid directoid devices across 59 microchannels), whereas straight channels show only a 3% difference in firing rate (*n* = 4 connectoid devices across 63 microchannels). Statistical significance between firing rates in the Q1 and Q4 regions for each microchannel was determined using a Mann-Whitney U test (*P* < 0.05). **(F)** Cross comparison of flux ratios plotted across 62 straight channels and 64 directional channels. Straight microchannels have ratios near 1, whereas directional channels approach is significantly larger with ratios near 3 (*P* = 1.67×10^−8^ Mann-Whitney U test).

We next evaluated the mean firing rate for all electrodes that fell within the first and last quartile distance along the length of each channel, 54% of directional microchannels had a significant difference in firing rate between first and last quartiles, whereas only 3% of straight microchannels demonstrated a significant difference in firing rate between the first and last quartiles (Fig. 5D,E). No significant increase was observed in straight microchannels (Firing rate data for each microstructure can be found in Supplementary Fig. 4). We observed significantly higher flux ratios in directional microchannels relative to straight microchannels (Fig 5F, *P* = 1.67×10^−8^, Mann-Whitney U test, straight microchannels = 64, 4 microchannels/condition/replicate, directional microchannels = 60 microchannels, 3 microstructures/condition replicate). Together, these results demonstrate a bioengineering strategy for monitoring directional action potential propagation between developing organoid models, with tunable connection weights defined by both the directionality and density of connectivity between spatially localized organoids.

## Discussion

Directionally weighted long-range projections are fundamental to the assembly of hierarchical brain circuits^6^, yet how these connections emerge and influence regional maturation and functional dynamics during early human development remains poorly understood. Here, we establish that stem cell-derived cortical and thalamic organoids can be assembled into a bioengineered system that preserves anatomical spatial identity while imposing directional bias on interregional connectivity we refer to as ‘directoids’. Using asymmetric microfluidic channels, we show that axon outgrowth can be reproducibly routed along permissive paths and strongly restricted in prohibitive ones, yielding structured asymmetry in long-range organoid-to-organoid projections. This engineered architecture is also expressed functionally, as directoids support directional action potential propagation and asymmetric firing-rate distributions not observed in straight-channel ‘connectoid’ controls. Together, these findings establish directoids as a scalable platform for reconstructing key features of developing interregional circuits and for probing how directional connectivity contributes to circuit assembly, regional specification, and functional integration in human neural tissues.

Our work builds on the emergence of ‘assembloids’ as a powerful strategy for modelling interactions between anatomically distinct brain regions in vitro^23,24,33–37^. These fused brain region specific systems have opened new opportunities to study neurodevelopmental transitions that are otherwise inaccessible *in vivo*, particularly those involving interaction between cortical and subcortical tissues^7^. Despite this promise, simple tissue fusions often lose anatomical organization over time because of cell migration across tissue boundaries, limiting their utility for studying long-range circuit organization^44^. To overcome this limitation, we designed and fabricated microfluidic chips that enable spatial localization of millimeter-scale organoids connected by micrometer-scale channels capable of directionally biasing long-range axonal projections (Fig. 2A). Our directoid microstructure facilitates axon growth in a permissive direction while blocking growth in the prohibitive direction. We validated this approach using live-cell imaging and with organoids engineered to express fluorescent proteins. Although our initial design iterations were inspired by prior demonstrations of growth cone control using asymmetric geometries^47–50^, we found that designs optimized for two-dimensional cultures were suboptimal for three-dimensional organoids, which generated excessive axonal input into the micrometer-scale channels (Supplementary Fig. 2). Our optimized design significantly reduced axon navigation in the prohibitive direction while promoting directional growth along permissive paths (Fig. 2B-D), producing significant differences in the total distance axons could traverse in forward versus reverse directions (Fig. 2E). In this way, our system advances beyond recent linear microchannel approaches for modeling reciprocal connectivity between organoids by enabling tunable control over long-range axonal connectivity^44^.

Long before the brain can process sensory input, hierarchically organized and reciprocally connected networks self-organize within and across the cortex and thalamus^18,67^. This process is driven internally, as progenitor cells and developing neurons are exposed to cascading waves of molecular cues^14,15^ and intrinsically generated neuronal activity^16,17,31,68^ that together establish the bidirectional functional circuits underlying the corticothalamic loop^19^. In our work, we build on these established protocols and to model region specific dorsal forebrain^62^ and thalamic^33^ organoids (Fig. 3A,B) that, when connected through our microfluidic platform, form regionally tunable, bidirectional long-range axonal projections, which can model asymmetric and bidirectional circuitry found in the corticothalamic loop *in vivo* (Fig. 4A)^6^. Our microfluidic design strategy enables rapid iteration with tunable control over spatial and directional axon targeting. We observed a 70.4% success rate of neurite transmission through permissive channel geometries to downstream organoid targets, compared to 0% in prohibitive channel geometry (Fig. 4B).

Finally, we confirmed that our directoid model can directionally route neuronal action potentials across anatomically distinct organoids using high-density CMOS-based microelectrode arrays. By analyzing spontaneous propagation events across microchannels, we quantified the efficacy of signal transmission through our channel designs. Asymmetric microchannels showed significantly higher firing rates at directional channel exits than at entrances, consistent with the presence of axons from both proximal and distal organoids at the exits. In contrast, straight channels displayed more uniform firing rates across microchannels and minimal variation in axon density along their lengths. Together, these results show that our microstructure interface is readily compatible with high-density electrophysiological readouts and can be used to model the directional assembly of developing circuits, providing a platform to investigate how cortical and subcortical networks process external inputs and extrinsic cues^69,70^.

Recent work in corticothalamic assembloids, has shown that Neuriexin1 (NRXN1)-mediated thalamic innervation of cortical organoid increases cortical cell diversity and expands the proportions of upper layer neurons, as reflected by increased expression of cortical markers such SATB2 and CTIP2^71^, supporting the *protocortex hypothesis*. In addition, reciprocal connectivity within corticothalamic assembloids has been shown to support synaptic plasticity, including both long-term potentiation and long-term depression^72^. These advances create new opportunities to investigate how tuned ratios of region-specific neuronal projections influence early development and functional maturation in controlled engineered environments.

Our demonstration of long-range axon guidance spanning hundreds of microns, without the use of molecular cues such as chemoattractants or chemorepellents^73^ establishes a reductionist organoid-based strategy for modeling brain region-specific circuit interactions^74,75^. In summary, we have developed a simple, scalable microfluidic platform based on repeating asymmetric geometries to direct the polarity of axonal projections between organoids. When combined with multimodal readouts such as calcium imaging^29,76^, genetically encoded neurotransmitter indicators^77,78^, and high-density electrophysiology readouts^30,55^, this system has the potential to provide and experimental path towards understanding how dynamic subcortical inputs shape cortical arealization^7^, and how human-specific genes govern self-assembly during early brain development^34,35,74,79^.

## Methods

### Fabrication of microfluidic device

Different devices were designed for different directional conditions. Lengths of channels varied between 700 μm to 2000 μm. Photo-patternable positive master molds were fabricated on silicon wafers in 3 layers consisting of copper alignment layer, then two successive SU-8 layers spin coated onto the wafer. The first SU-8-2 (Kayaku, Cat. Y131240 0500L1GL) layer formed directional channels at a height of 3.5 μm. The second layer of SU-8-100 (Kayaku, Cat. Y131240 0500L1GL) formed the culture compartment at 300 μm. Master molds were silanized (Sigma Aldrich, Cat. Y131240 0500L1GL) in a vacuum chamber for 20 minutes, then cured at 65°C for 120 minutes. Devices were replicated by mixing polydimethylsiloxane and curing agent 1:10 (PDMS) (Sylguard 184, Cat. 761036), then degassing for 20 minutes.

PDMS was spin coated to a height of 200 μm and cured at 65°C for 4 hours. Uncured PDMS was removed after curing by soaking microstructures in ethyl acetate in a crystallization dish for 12 hours, then washed rapidly for 5 minutes in 95% isopropyl alcohol (IPA) and dried with N2. Microstructures and glass coverslips were autoclaved prior to attachment to fully sterilize all surfaces.

PDMS microstructures were either attached to autoclaved coverslips cleaned with 37% hydrochloric acid (HCl) (Sigma Alrich, Cat. Y131240 0500L1GL) overnight then washed with deionized water 3 times, then IPA. For long term imaging, microstructures were attached to glass bottom tissue culture plates. Plates or coverslips are initially treated with 0.07% Poly(ethyleneimine) (PEI) solution, washed 3x times in tissue culture grade water, then fully dried via aspirating pipette. Microstructures are then attached to glass coverslip. 50 ug/mL Laminin (Cat: L2020-1MG, Sigma-Alrich) is dispensed on both well ends, then placed under vacuum at room temperature for 20 minutes to ensure full coating of inner wells. Channel openness and hydrophilicity is evaluated on one sacrificial microstructure by adding 1:100 dilution of 2 µm polystyrene beads (Cat: L3030, Sigma-Alrich), desiccated under vacuum for an additional 20 minutes, then checked under a fluorescence microscope.

### Cell Culture

#### Human organoid generation

Two fluorescently labelled human embryonic stem cell (hESC) lines used (H9-cag-mCherry, WiCell, Cat. WB67905 and H9-CreLoxP-GFP, WiCell. Cat. WB20971) were used to generate brain region specific organoids of dorsal forebrain and dorsal thalamus identity following established protocol base on name Yoon *et al*^62^ and Xiang *et al*.^26^. hESC lines were thawed with 5 µM Rock Inhibitor into 6 well plates coated with 1.7% hESC qualified Matrigel and maintained in mTesr supplemented with 10ng/mL FGF-2 media with daily media exchanges to maintain pluripotency. Aggregation occurred when hESC plates reached 70-80% confluency. hESCs were lifted by removing media and adding 1mL TryplE Select for 6 minutes at 37°C, then using a P1000 to manually triturate cells added to 5mL of warmed mTesr media to dilute in, then spun down for 3 minutes at 200RCF, resuspended in 1mL of media and counted for organoid aggregation.

#### Human dorsal cortex organoid generation

Dorsal cortex spheroids were generated when H9-mCherry cells were seeded at 1×10^5^ cells/well in 96 well slit-well plates and maintained in these plates for the duration of culturing. Dorsal organoids were generated after the method of Yoon et al^62^ with modifications.^62^ with modifications. Day 1-6 media was changed daily with dorsal cortex neural induction media (DCM1), consisting of E6 base media with 1% Pen/Strep and at time of feeding was supplemented with small molecules 10µM SB-431542, 0.15 µM LDN, and 2.5 µM XAV-939.

From day 7 to day 25, dorsal cortex neural patterning media (DCM2) consisting of Neurobasal base medium supplemented with 2% B-27 (without Vitamin A), 1% Glutamax, and 1% Pen/Strep. From day 19 to day 24, DCM2 was supplemented with 20 µM FGF-2 and 20 µM EGF-1 added immediately before feeding. DCM2 media was exchanged every other day.

From day 26 to day 43, dorsal cortex maturation media (DCM3) was exchanged every 3-4 days. DCM3 consists of neurobasal base media supplemented with 2% B-27 (with Vitamin A), 1% Glutamax, and 1% Pen/Strep. Neurotrophic growth factors consisting of 20ng/mL NT3 and 20 ng/mL BDNF added to DCM3 immediately before feeding.

#### Human dorsal thalamus organoid generation

Dorsal Thalamus spheroids were generated when H9-creLox-GFP hESCs (H9-CreLox-GFP, WiCell. Cat. WB20971) were seeded at 9×10^3^ cells/well into 96 well slit well plates and maintained in these plates for the duration of culturing. Thalamic organoids were generated after the method of Xiang *et al*.^26^ with modifications.

From days 1 to 8 media was changed daily with Thalamic Induction Media (Th1), consisting of DMEM/F12 basal media supplemented with 15% Knockout Serum Replacement (KSR), 1% Glutamax, 1% MEM-NEAA, 100 µm β-mercaptoethanol (BME) and 50 ug/mL Primocin. On aggregation day (Day 0), Th1 supplemented with 100 nM LDN, 10 uM SB-431542, 5% heat inactivated FBS, 4ug/mL Insulin and 50uM Rock Inhibitor added immediately before media exchange. On Day 2, Th1 was exchanged with small molecules 100 nM LDN, 10 uM SB-431542, 4ug/mL Insulin and 50uM Rock Inhibitor immediately before feeding. On days 4 and 6, Th1 was exchanged with small molecules 100 nM LDN, 10 uM SB-431542, and 4ug/mL Insulin added immediately before feeding.

From day 8 to day 16, Thalamic Patterning Media (Th2) was changed every other day, consisting of DMEM/F12 basal media supplemented with 0.15% of Dextrose, 100 µM BME, 1% N2, 2% B27 (without Vitamin A) and 50ug/mL Primocin. Small molecules 30 ng/mL BMP7 and 1 µM PD327901 added immediately before feeding.

From Day 16 on, Thalamic Maturation Media (Th3) was exchanged every other day until Day 25, then fed every 3 days thereafter. Th3 media consists of a 50:50 mix of DMEM/F12: Neurobasal base media was supplemented with 0.5% N2, 1% B27 (with Vitamin A), 1% Glutamax, 0.5% MEM-NEAA, 50 µM BME, 2.6ug/mL Insulin, and 50ug/mL Primocin.

Neurotrophic Growth factors consisting of 200 µM Ascorbic Acid and 20 ng/mL BDNF added immediately before feeding.

### Mouse Thalamic Organoid Generation

Dorsal Thalamus spheroids were generated from Bruce4 mESCs (Bruce4-C57/BL6 Millipore. Cat. CMTI-2) using methods adapted from Kiral-Park *et al*^26^ were seeded at 3×10^3^ cells/well into 96 well slit well plates and maintained in these plates for the duration of culturing. Thalamic organoids were generated after the method of Kiral et al^62^ with modifications. From days 1 to 8 media was changed daily with Thalamic Induction Media (mTh1), consisting of DMEM/F12 basal media supplemented with 15% Knockout Serum Replacement (KSR), 1% Glutamax, 1% MEM-NEAA, 100 µm β-mercaptoethanol (BME) and 50 ug/mL Primocin. On aggregation day (Day 0), mTh1 supplemented with 100 nM LDN, 10 uM SB-431542, 5% heat inactivated FBS, 4ug/mL Insulin and 50uM Rock Inhibitor added immediately before media exchange. On Day 2, mTh1 was exchanged with small molecules 100 nM LDN, 10 uM SB-431542, 4ug/mL Insulin and 50uM Rock Inhibitor immediately before feeding. On days 4 and 6, mTh1 was exchanged with small molecules 100 nM LDN, 10 uM SB-431542, and 4ug/mL Insulin added immediately before feeding.

From day 7 to day 15, Thalamic Patterning Media (mTh2) was changed every other day, consisting of DMEM/F12 basal media supplemented with 0.15% of Dextrose, 100 µM BME, 1% N2, 2% B27 (without Vitamin A) and 50ug/mL Primocin. Small molecules 30 ng/mL BMP7 and 1 µM PD327901 added immediately before feeding.

From Day 16 on, Thalamic Maturation Media (mTh3) was exchanged every other day until Day 25, then fed every 3 days thereafter. Th3 media consists of a BrainPhys base media supplemented with 0.5% N2, 1% B27 (with Vitamin A), 10ug/mL Heparin, 1% Glutamax, 0.5% MEM-NEAA, and 50ug/mL Primocin. 200 µM Ascorbic Acid was added immediately before feeding.

### Slice Organoid Preparation

For slice organoids, 30 to 35-day old organoids were taken out of the incubator and transferred to sterile 1.5mL Eppendorf tubes with 1mL of media. Organoids were allowed to equilibrate for 5 minutes, then placed in 4 °C and allowed to cool for an additional 10 minutes. Organoids were then embedded in 4% agarose reconstituted in DMEM/F12 media with 4% Pen/Strep to minimize contamination and sliced at 300 µm using a vibratome. Slices were removed to a 24 well plate with 4% Pen/Strep, allowed to incubate a minimum of 5 minutes in the biosafety cabinet, then transferred to successive wells containing DMEM/F12 at 3%, 2%, and 2% Pen/Strep for 5 minutes each to minimize contamination before plating.

### Organoid Seeding

Organoids were plated in microstructure wells by pipetting organoids in, removing all media then adding a drop of hydrogel (Geltrex, Thermofisher) in the cell plating area to aid in cell attachment. Microstructures were then incubated for 10 minutes at 37°C, 5% CO_2_ to stabilize the hydrogel. Warmed media was then added to microstructures. Media was changed every other day with 2% Pen/Strep added to minimize contamination risk from the plating process.

### Live Imaging

Live imaging was performed by tracking axonal outgrowth three days after organoid seeding to confirm overall organoid health, firm cell attachment, and initial neurite outgrowth into the microchannels. Microstructures were imaged hourly for four days on a BioTek Lionheart FX Automated Microscope, with breaks every two days for media changes. The automated imaging system was maintained at 37°C, 5% humidity, with diH2O used to replenish the humidity reservoir surrounding the plate during media change intervals and restarting automated imaging.

### Immunohistochemistry and Imaging

Organoids were fixed in 4% PFA in Dubecco’s phosphate buffered saline (dPBS) at DIV 30 to DIV35 and DIV 55 to verify region specific patterning of human dorsal cortex and human thalamus. Primary antibodies used to identify neuronal markers for both types of organoids were SOX2, (1:250 Mouse, Cat. sc-365823, Santa Cruz Biotechnologies), and MAP2 (1:3000, Mouse Cat.GTX82661, GeneTex). Primary antibodies used to confirm dorsal forebrain identity were FOXG1 (1:100 Rabbit, Cat. NBP3-04084, R&D Systems), CTIP2 (1:200 Rat, Cat. ab18465, Abcam). Primary antibodies used to identify thalamic identity were FOXP2 (1:250 Mouse, Cat. sc-517261, Santa Cruz Biotechnologies), TUJ1 (1:2000 Mouse, Cat. 801201, Bioloegend), and PAX6 (1:100 Mouse, Cat. ab195045, Abcam). Because H9 lines were either CAG-mCherry or creLox-GFP tagged, secondary antibodies for our markers of interest were selected to fluoresce green (Alexa Fluor 546: Goat α-Mouse IgG Cat. ab150113, AbCam/Goat α-Chicken IgY, Cat. ab150173 Abcam/Donkey αRabbit IgG Cat. ab150073, AbCam) to visualize markers for mCherry lines and Red (Alexa Fluor 568 Cat.ab175471, AbCam) or FarRed (AlexaFluor 647, Cat. ab150135, AbCam) for GFP cell lines. Nuclei were stained with Hoescht dye (1:10,000, Cat. H1399, Life Technologies), incubated for 20 minutes at room temperature, then washed twice with dPBS before imaging. Organoids were mounted on coverslips then imaged under either EVOS FL fluorescent microscope or Zeiss 880 confocal microscope to image FarRed tagged regions.

### MEA microstructure chip preparation

Recordings were performed on planar surface of high-density CMOS-based microelectrode arrays containing 26,400 recording sites, with 1,024 sites available for simultaneous recording^55^.^55^. The approximately 8 mm² planar recording area bonds to PDMS microstructures with Van der Waal forces and have a cross-sectional footprint that that can resolve microfluidic attachment (**Supplementary Fig.3**). High density microelectrode array (MEAs)MaxOne+ chips from MaxWell Biosystems were sanitized with 70% ethanol, Terg-y-Zyme for 1 hour, rinsed 3x times with diH2O, then blow dried with N2. Sterile microstructures were attached to MEA surface with autoclaved forceps and adhered with electrostatic forces. PBS was then added to the microchip and placed in a vacuum desiccator to remove bubbles in the channels. Channel openness was confirmed via a custom Python script that generated impedance mapping of PDMS over the electrodes, (provided by Jens Duru). 50 ug/mL Laminin (Cat: L2020-1MG, Sigma-Alrich) is dispensed on both well ends, then placed under vacuum at room temperature for 20 minutes to ensure full coating of inner wells. Only thalamic organoids were used in this experiment. All media was removed via pipette and a drop of Geltrex was added to aid in cell attachment. Microstructures were then incubated for 10 minutes at 37°C, 5% CO_2_ to stabilize hydrogel. Warmed media was then added to microstructures. Media was changed every other day with 2% Pen/Strep added to minimize contamination risk from the plating process. Custom electrode sampling configurations was optimized to capture microstructure channels by overlaying each corresponding impedance map on their respective chips and selecting channel regions only. 10 minute electrophysiology recordings of channels were taken 10 days post seeding.

### Electrophysiology Analysis

Peak detection was performed on raw H5 recording output files with SciPy peaks at >3.5 RMS with ISI removal <1ms and was performed via a custom Python script. To generate latency trace plots, the impedance map was overlaid to manually pairs of electrodes at entry (Source) and exit (Target) respectively for a single microchannel. If a spike occurred initially at the Source electrode at *t* = 0 and a co-occurring spike was also observed at the Target electrode with *t*=1-2 ms latency, these paired spikes were defined as a potential propagation event. A minimum of 30 potential propagation events were required for propagation analysis of all selected pairs. A principal axis was defined between the Source/Target electrodes and a surrounding 17.5 μm (1 electrode adjacent) orthogonal search band filtered for possible intermediary electrode candidates. For all paired spike time windows that occurred for Source/Target electrodes, spike counts for intermediary candidates were printed as well as their relative positions along the principal axis. Intermediary candidates were selected at approximately 25%, 50%, and 75% distances along the principal axis if corresponding co-occurring spikes were also observed in more than 80% of the paired spike windows.

For each quartile, a Source and Target electrode was selected to maintain consistent sampling for each microchannel in the latency trace plots. All intermediate electrodes were grouped by interquartile distance along the principal axis between the source and target. For each group of electrodes, we aggregated spike counts, then normalized all groups to firing rate (Hz) by dividing the # of peaks detected/recording duration for each electrode sampled. ISI violating peaks were removed from analysis.

Spike frequency ratio was defined as the ratio of the average spiking frequency for electrodes sampled at FR(bin_n__/FR(25%). For Fig. 5D, 5% bins were selected for sampling ranges to illustrate changes in firing rates along a microchannel. For **Fig. 5E** and F, the first 25% and last 25% average firing rate were compared. MannWhitney-U test was performed in Fig. 5E for all sampled electrodes in the first and last quartile of a single selected channel to determine significance for both straight and asymmetric channels. Firing rate boxplots for microstructures and their channels are in **Supplementary Fig. 5.5**.

## Code Availability

Analysis was performed with custom Python scripts which are publicly available at: https://github.com/MightyMolecule/spikePathTool

Impedance mapping was performed with scripts from J.Duru, is publicly available at: https://github.com/lbb-neuron/CMOS_stimulation_and_recording

## Acknowledgements

The authors would like to thank members of the Braingeneers consortium for helpful discussions. This research was supported by the National Institutes of Health T32 NIH Award (5T32HG012344-04 to A.C.). The NIH IRADCA also supported this research Award (K12GM139185 to M.M.). This study was supported by the National Science Foundation (NSF) Emerging Frontiers in Research and Innovation under award (NSF 2515389 to T.S. and M.T.) and the Brain and Behavior Research Foundation Young Investigator Award (to T.S), and the NIH Outstanding Investigator Award (R35GM147395-04 to A.S.). The authors acknowledge technical support from Benjamin Abrams, UCSC Life Sciences Microscopy Center, RRID: SCR_021135.

## Author contributions

T.S. conceived and supervised the study. A.C. designed, fabricated microstructures, performed cell work and imaging, made electrophysiology recordings, wrote scripts and performed computational analysis and statistics for this study. M.M designed, fabricated microstructures, performed cell work, immunohistochemistry, and imaging for this study. K.K., T.G., Z.G., A.Z., and SM performed immunohistochemistry for this study. A.A. provided technical support on fabrication development. N.W. assisted with figure formatting and code review. G.K provided assembloid images for the graphical abstract. E.S. provided technical imaging support. A.C. wrote the first draft of the manuscript, T.S. provided subsequent edits, and all authors discussed the results and commented on the paper.

## References

(1) Nikolopoulou, E.; Galea, G. L.; Rolo, A.; Greene, N. D. E.; Copp, A. J. Neural Tube Closure: Cellular, Molecular and Biomechanical Mechanisms. Development 2017, 144 (4), 552–566. 10.1242/dev.145904.

(2) Xue, X.; Kim, Y. S.; Ponce-Arias, A.-I.; O’Laughlin, R.; Yan, R. Z.; Kobayashi, N.; Tshuva, R. Y.; Tsai, Y.-H.; Sun, S.; Zheng, Y.; Liu, Y.; Wong, F. C. K.; Surani, A.; Spence, J. R.; Song, H.; Ming, G.-L.; Reiner, O.; Fu, J. A Patterned Human Neural Tube Model Using Microfluidic Gradients. Nature 2024, 628 (8007), 391–399. 10.1038/s41586-024-07204-7.

(3) Karzbrun, E.; Khankhel, A. H.; Megale, H. C.; Glasauer, S. M. K.; Wyle, Y.; Britton, G.; Warmflash, A.; Kosik, K. S.; Siggia, E. D.; Shraiman, B. I.; Streichan, S. J. Human Neural Tube Morphogenesis in Vitro by Geometric Constraints. Nature 2021, 599 (7884), 268–272. 10.1038/s41586-021-04026-9.

(4) Assaf, Y.; Bouznach, A.; Zomet, O.; Marom, A.; Yovel, Y. Conservation of Brain Connectivity and Wiring across the Mammalian Class. Nat. Neurosci. 2020, 23 (7), 805–808. 10.1038/s41593-020-0641-7.

(5) Bullmore, E.; Sporns, O. Complex Brain Networks: Graph Theoretical Analysis of Structural and Functional Systems. Nat. Rev. Neurosci. 2009, 10 (3), 186–198. 10.1038/nrn2575.

(6) Oh, S. W.; Harris, J. A.; Ng, L.; Winslow, B.; Cain, N.; Mihalas, S.; Wang, Q.; Lau, C.; Kuan, L.; Henry, A. M.; Mortrud, M. T.; Ouellette, B.; Nguyen, T. N.; Sorensen, S. A.; Slaughterbeck, C. R.; Wakeman, W.; Li, Y.; Feng, D.; Ho, A.; Nicholas, E.; Hirokawa, K. E.; Bohn, P.; Joines, K. M.; Peng, H.; Hawrylycz, M. J.; Phillips, J. W.; Hohmann, J. G.; Wohnoutka, P.; Gerfen, C. R.; Koch, C.; Bernard, A.; Dang, C.; Jones, A. R.; Zeng, H. A Mesoscale Connectome of the Mouse Brain. Nature 2014, 508 (7495), 207–214. 10.1038/nature13186.

(7) Cadwell, C. R.; Bhaduri, A.; Mostajo-Radji, M. A.; Keefe, M. G.; Nowakowski, T. J. Development and Arealization of the Cerebral Cortex. Neuron 2019, 103 (6), 980–1004. 10.1016/j.neuron.2019.07.009.

(8) Bullmore, E.; Sporns, O. The Economy of Brain Network Organization. Nat. Rev. Neurosci. 2012, 13 (5), 336–349. 10.1038/nrn3214.

(9) Ma, Z.; Turrigiano, G. G.; Wessel, R.; Hengen, K. B. Cortical Circuit Dynamics Are Homeostatically Tuned to Criticality In Vivo. Neuron 2019, 104 (4), 655–664.e4. 10.1016/j.neuron.2019.08.031.

(10) Nano, P. R.; Fazzari, E.; Azizad, D.; Martija, A.; Nguyen, C. V.; Wang, S.; Giang, V.; Kan, R. L.; Yoo, J.; Wick, B.; Haeussler, M.; Bhaduri, A. Integrated Analysis of Molecular Atlases Unveils Modules Driving Developmental Cell Subtype Specification in the Human Cortex. Nat. Neurosci. 2025, 1–15. 10.1038/s41593-025-01933-2.

(11) Tasic, B.; Yao, Z.; Graybuck, L. T.; Smith, K. A.; Nguyen, T. N.; Bertagnolli, D.; Goldy, J.; Garren, E.; Economo, M. N.; Viswanathan, S.; Penn, O.; Bakken, T.; Menon, V.; Miller, J.; Fong, O.; Hirokawa, K. E.; Lathia, K.; Rimorin, C.; Tieu, M.; Larsen, R.; Casper, T.; Barkan, E.; Kroll, M.; Parry, S.; Shapovalova, N. V.; Hirschstein, D.; Pendergraft, J.; Sullivan, H. A.; Kim, T. K.; Szafer, A.; Dee, N.; Groblewski, P.; Wickersham, I.; Cetin, A.; Harris, J. A.; Levi, B. P.; Sunkin, S. M.; Madisen, L.; Daigle, T. L.; Looger, L.; Bernard, A.; Phillips, J.; Lein, E.; Hawrylycz, M.; Svoboda, K.; Jones, A. R.; Koch, C.; Zeng, H. Shared and Distinct Transcriptomic Cell Types across Neocortical Areas. Nature 2018, 563 (7729), 72–78. 10.1038/s41586-018-0654-5.

(12) Bishop, K. M.; Goudreau, G.; O’Leary, D. D. M. Regulation of Area Identity in the Mammalian Neocortex by Emx2 and Pax6. Science 2000, 288 (5464), 344–349. 10.1126/science.288.5464.344.

(13) Rakic, P. Specification of Cerebral Cortical Areas. Science 1988, 241 (4862), 170–176. 10.1126/science.3291116.

(14) Creutzfeldt, O. D. Generality of the Functional Structure of the Neocortex. Naturwissenschaften 1977, 64 (10), 507–517. 10.1007/BF00483547.

(15) O’Leary, D. D. M. Do Cortical Areas Emerge from a Protocortex? Trends Neurosci. 1989, 12 (10), 400–406. 10.1016/0166-2236(89)90080-5.

(16) Ackman, J. B.; Burbridge, T. J.; Crair, M. C. Retinal Waves Coordinate Patterned Activity throughout the Developing Visual System. Nature 2012, 490 (7419), 219–225. 10.1038/nature11529.

(17) Chini, M.; Pfeffer, T.; Hanganu-Opatz, I. An Increase of Inhibition Drives the Developmental Decorrelation of Neural Activity. eLife 2022, 11, e78811. 10.7554/eLife.78811.

(18) Kim, C. N.; Shin, D.; Wang, A.; Nowakowski, T. J. Spatiotemporal Molecular Dynamics of the Developing Human Thalamus. Science 2023, 382 (6667), eadf9941. 10.1126/science.adf9941.

(19) Shepherd, G. M. G.; Yamawaki, N. Untangling the Cortico-Thalamo-Cortical Loop: Cellular Pieces of a Knotty Circuit Puzzle. Nat. Rev. Neurosci. 2021, 22 (7), 389–406. 10.1038/s41583-021-00459-3.

(20) Eiraku, M.; Watanabe, K.; Matsuo-Takasaki, M.; Kawada, M.; Yonemura, S.; Matsumura, M.; Wataya, T.; Nishiyama, A.; Muguruma, K.; Sasai, Y. Self-Organized Formation of Polarized Cortical Tissues from ESCs and Its Active Manipulation by Extrinsic Signals. Cell Stem Cell 2008, 3 (5), 519–532. 10.1016/j.stem.2008.09.002.

(21) Giandomenico, S. L.; Mierau, S. B.; Gibbons, G. M.; Wenger, L. M. D.; Masullo, L.; Sit, T.; Sutcliffe, M.; Boulanger, J.; Tripodi, M.; Derivery, E.; Paulsen, O.; Lakatos, A.; Lancaster, M. A. Cerebral Organoids at the Air–Liquid Interface Generate Diverse Nerve Tracts with Functional Output. Nat. Neurosci. 2019, 22 (4), 669–679. 10.1038/s41593-019-0350-2.

(22) Lancaster, M. A.; Renner, M.; Martin, C.-A.; Wenzel, D.; Bicknell, L. S.; Hurles, M. E.; Homfray, T.; Penninger, J. M.; Jackson, A. P.; Knoblich, J. A. Cerebral Organoids Model Human Brain Development and Microcephaly. Nature 2013, 501 (7467), 373–379. 10.1038/nature12517.

(23) Birey, F.; Andersen, J.; Makinson, C. D.; Islam, S.; Wei, W.; Huber, N.; Fan, H. C.; Metzler, K. R. C.; Panagiotakos, G.; Thom, N.; O’Rourke, N. A.; Steinmetz, L. M.; Bernstein, J. A.; Hallmayer, J.; Huguenard, J. R.; Paşca, S. P. Assembly of Functionally Integrated Human Forebrain Spheroids. Nature 2017, 545 (7652), 54–59. 10.1038/nature22330.

(24) Wang, Y.; Chiola, S.; Yang, G.; Russell, C.; Armstrong, C. J.; Wu, Y.; Spampanato, J.; Tarboton, P.; Ullah, H. M. A.; Edgar, N. U.; Chang, A. N.; Harmin, D. A.; Bocchi, V. D.; Vezzoli, E.; Besusso, D.; Cui, J.; Cattaneo, E.; Kubanek, J.; Shcheglovitov, A. Modeling Human Telencephalic Development and Autism-Associated SHANK3 Deficiency Using Organoids Generated from Single Neural Rosettes. Nat. Commun. 2022, 13 (1), 5688. 10.1038/s41467-022-33364-z.

(25) Gordon, A.; Yoon, S.-J.; Tran, S. S.; Makinson, C. D.; Park, J. Y.; Andersen, J.; Valencia, A. M.; Horvath, S.; Xiao, X.; Huguenard, J. R.; Pașca, S. P.; Geschwind, D. H. Long-Term Maturation of Human Cortical Organoids Matches Key Early Postnatal Transitions. Nat. Neurosci. 2021, 24 (3), 331–342. 10.1038/s41593-021-00802-y.

(26) Kiral, F. R.; Cakir, B.; Tanaka, Y.; Kim, J.; Yang, W. S.; Wehbe, F.; Kang, Y.-J.; Zhong, M.; Sancer, G.; Lee, S.-H.; Xiang, Y.; Park, I.-H. Generation of Ventralized Human Thalamic Organoids with Thalamic Reticular Nucleus. Cell Stem Cell 2023, 30 (5), 677–688.e5. 10.1016/j.stem.2023.03.007.

(27) Quadrato, G.; Nguyen, T.; Macosko, E. Z.; Sherwood, J. L.; Min Yang, S.; Berger, D. R.; Maria, N.; Scholvin, J.; Goldman, M.; Kinney, J. P.; Boyden, E. S.; Lichtman, J. W.; Williams, Z. M.; McCarroll, S. A.; Arlotta, P. Cell Diversity and Network Dynamics in Photosensitive Human Brain Organoids. Nature 2017, 545 (7652), 48–53. 10.1038/nature22047.

(28) Fair, S. R.; Julian, D.; Hartlaub, A. M.; Pusuluri, S. T.; Malik, G.; Summerfied, T. L.; Zhao, G.; Hester, A. B.; Ackerman, W. E.; Hollingsworth, E. W.; Ali, M.; McElroy, C. A.; Buhimschi, I. A.; Imitola, J.; Maitre, N. L.; Bedrosian, T. A.; Hester, M. E. Electrophysiological Maturation of Cerebral Organoids Correlates with Dynamic Morphological and Cellular Development. Stem Cell Rep. 2020, 15 (4), 855–868. 10.1016/j.stemcr.2020.08.017.

(29) Samarasinghe, R. A.; Miranda, O. A.; Buth, J. E.; Mitchell, S.; Ferando, I.; Watanabe, M.; Allison, T. F.; Kurdian, A.; Fotion, N. N.; Gandal, M. J.; Golshani, P.; Plath, K.; Lowry, W. E.; Parent, J. M.; Mody, I.; Novitch, B. G. Identification of Neural Oscillations and Epileptiform Changes in Human Brain Organoids. Nat. Neurosci. 2021, 24 (10), 1488–1500. 10.1038/s41593-021-00906-5.

(30) Sharf, T.; van der Molen, T.; Glasauer, S. M. K.; Guzman, E.; Buccino, A. P.; Luna, G.; Cheng, Z.; Audouard, M.; Ranasinghe, K. G.; Kudo, K.; Nagarajan, S. S.; Tovar, K. R.; Petzold, L. R.; Hierlemann, A.; Hansma, P. K.; Kosik, K. S. Functional Neuronal Circuitry and Oscillatory Dynamics in Human Brain Organoids. Nat. Commun. 2022, 13 (1), 4403. 10.1038/s41467-022-32115-4.

(31) Molen, T. van der; Spaeth, A.; Chini, M.; Hernandez, S.; Kaurala, G. A.; Schweiger, H. E.; Duncan, C.; McKenna, S.; Geng, J.; Lim, M.; Bartram, J.; Dendukuri, A.; Zhang, Z.; Gonzalez-Ferrer, J.; Bhaskaran-Nair, K.; Blauvelt, L. J.; Harder, C. R. K.; Petzold, L. R.; Din, D.-M. A. E.; Laird, J.; Schenke, M.; Smirnova, L.; Colquitt, B. M.; Mostajo-Radji, M. A.; Hansma, P. K.; Teodorescu, M.; Hierlemann, A.; Hengen, K. B.; Hanganu-Opatz, I. L.; Kosik, K. S.; Sharf, T. Protosequences in Brain Organoids Model Intrinsic Brain States. bioRxiv January 8, 2025, p 2023.12.29.573646. 10.1101/2023.12.29.573646.

(32) Miura, Y.; Li, M.-Y.; Birey, F.; Ikeda, K.; Revah, O.; Thete, M. V.; Park, J.-Y.; Puno, A.; Lee, S. H.; Porteus, M. H.; Pașca, S. P. Generation of Human Striatal Organoids and Cortico-Striatal Assembloids from Human Pluripotent Stem Cells. Nat. Biotechnol. 2020, 38 (12), 1421–1430. 10.1038/s41587-020-00763-w.

(33) Xiang, Y.; Tanaka, Y.; Cakir, B.; Patterson, B.; Kim, K.-Y.; Sun, P.; Kang, Y.-J.; Zhong, M.; Liu, X.; Patra, P.; Lee, S.-H.; Weissman, S. M.; Park, I.-H. hESC-Derived Thalamic Organoids Form Reciprocal Projections When Fused with Cortical Organoids. Cell Stem Cell 2019, 24 (3), 487–497.e7. 10.1016/j.stem.2018.12.015.

(34) Shin, D.; Kim, C. N.; Ross, J.; Hennick, K. M.; Wu, S.-R.; Paranjape, N.; Leonard, R.; Wang, J. C.; Keefe, M. G.; Pavlovic, B. J.; Donohue, K. C.; Moreau, C.; Wigdor, E. M.; Larson, H. H.; Allen, D. E.; Cadwell, C. R.; Bhaduri, A.; Popova, G.; Bearden, C. E.; Pollen, A. A.; Jacquemont, S.; Sanders, S. J.; Haussler, D.; Wiita, A. P.; Frost, N. A.; Sohal, V. S.; Nowakowski, T. J. Thalamocortical Organoids Enable in Vitro Modeling of 22q11.2 Microdeletion Associated with Neuropsychiatric Disorders. Cell Stem Cell 2024, 31 (3), 421–432.e8. 10.1016/j.stem.2024.01.010.

(35) Wu, S.-R.; Nowakowski, T. J. Exploring Human Brain Development and Disease Using Assembloids. Neuron 2025, 113 (8), 1133–1150. 10.1016/j.neuron.2025.02.010.

(36) Bagley, J. A.; Reumann, D.; Bian, S.; Lévi-Strauss, J.; Knoblich, J. A. Fused Cerebral Organoids Model Interactions between Brain Regions. Nat. Methods 2017, 14 (7), 743–751. 10.1038/nmeth.4304.

(37) Kim, J.; Miura, Y.; Li, M.-Y.; Revah, O.; Selvaraj, S.; Birey, F.; Meng, X.; Thete, M. V.; Pavlov, S. D.; Andersen, J.; Pașca, A. M.; Porteus, M. H.; Huguenard, J. R.; Pașca, S. P. Human Assembloids Reveal the Consequences of *CACNA1G* Gene Variants in the Thalamocortical Pathway. Neuron 2024, 112 (24), 4048–4059.e7. 10.1016/j.neuron.2024.09.020.

(38) Zhu, Y.; Zhang, X.; Sun, L.; Wang, Y.; Zhao, Y. Engineering Human Brain Assembloids by Microfluidics. Adv. Mater. 2023, 35 (14), 2210083. 10.1002/adma.202210083.

(39) Reumann, D.; Krauditsch, C.; Novatchkova, M.; Sozzi, E.; Wong, S. N.; Zabolocki, M.; Priouret, M.; Doleschall, B.; Ritzau-Reid, K. I.; Piber, M.; Morassut, I.; Fieseler, C.; Fiorenzano, A.; Stevens, M. M.; Zimmer, M.; Bardy, C.; Parmar, M.; Knoblich, J. A. In Vitro Modeling of the Human Dopaminergic System Using Spatially Arranged Ventral Midbrain–Striatum–Cortex Assembloids. Nat. Methods 2023, 1–14. 10.1038/s41592-023-02080-x.

(40) O’Laughlin, R.; Cheng, F.; Song, H.; Ming, G. Bioengineering Tools for Next-Generation Neural Organoids. Curr. Opin. Neurobiol. 2025, 92, 103011. 10.1016/j.conb.2025.103011.

(41) Ao, Z.; Cai, H.; Wu, Z.; Ott, J.; Wang, H.; Mackie, K.; Guo, F. Controllable Fusion of Human Brain Organoids Using Acoustofluidics. Lab. Chip 2021, 21 (4), 688–699. 10.1039/D0LC01141J.

(42) Roth, J. G.; Huang, M. S.; Li, T. L.; Feig, V. R.; Jiang, Y.; Cui, B.; Greely, H. T.; Bao, Z.; Paşca, S. P.; Heilshorn, S. C. Advancing Models of Neural Development with Biomaterials. Nat. Rev. Neurosci. 2021, 22 (10), 593–615. 10.1038/s41583-021-00496-y.

(43) Cho, A.-N.; Jin, Y.; An, Y.; Kim, J.; Choi, Y. S.; Lee, J. S.; Kim, J.; Choi, W.-Y.; Koo, D.-J.; Yu, W.; Chang, G.-E.; Kim, D.-Y.; Jo, S.-H.; Kim, J.; Kim, S.-Y.; Kim, Y.-G.; Kim, J. Y.; Choi, N.; Cheong, E.; Kim, Y.-J.; Je, H. S.; Kang, H.-C.; Cho, S.-W. Microfluidic Device with Brain Extracellular Matrix Promotes Structural and Functional Maturation of Human Brain Organoids. Nat. Commun. 2021, 12 (1), 4730. 10.1038/s41467-021-24775-5.

(44) Osaki, T.; Duenki, T.; Chow, S. Y. A.; Ikegami, Y.; Beaubois, R.; Levi, T.; Nakagawa-Tamagawa, N.; Hirano, Y.; Ikeuchi, Y. Complex Activity and Short-Term Plasticity of Human Cerebral Organoids Reciprocally Connected with Axons. Nat. Commun. 2024, 15 (1), 2945. 10.1038/s41467-024-46787-7.

(45) Martins-Costa, C.; Wiegers, A.; Pham, V. A.; Sidhaye, J.; Doleschall, B.; Novatchkova, M.; Lendl, T.; Piber, M.; Peer, A.; Möseneder, P.; Stuempflen, M.; Chow, S. Y. A.; Seidl, R.; Prayer, D.; Höftberger, R.; Kasprian, G.; Ikeuchi, Y.; Corsini, N. S.; Knoblich, J. A. ARID1B Controls Transcriptional Programs of Axon Projection in an Organoid Model of the Human Corpus Callosum. Cell Stem Cell 2024, 31 (6), 866–885.e14. 10.1016/j.stem.2024.04.014.

(46) Taylor, A. M.; Rhee, S. W.; Tu, C. H.; Cribbs, D. H.; Cotman, C. W.; Jeon, N. L. Microfluidic Multicompartment Device for Neuroscience Research. Langmuir 2003, 19 (5), 1551–1556. 10.1021/la026417v.

(47) Peyrin, J.-M.; Deleglise, B.; Saias, L.; Vignes, M.; Gougis, P.; Magnifico, S.; Betuing, S.; Pietri, M.; Caboche, J.; Vanhoutte, P.; Viovy, J.-L.; Brugg, B. Axon Diodes for the Reconstruction of Oriented Neuronal Networks in Microfluidic Chambers. Lab. Chip 2011, 11 (21), 3663–3673. 10.1039/C1LC20014C.

(48) Renault, R.; Durand, J.-B.; Viovy, J.-L.; Villard, C. Asymmetric Axonal Edge Guidance: A New Paradigm for Building Oriented Neuronal Networks. Lab. Chip 2016, 16 (12), 2188–2191. 10.1039/C6LC00479B.

(49) Holloway, P. M.; Hallinan, G. I.; Hegde, M.; Lane, S. I. R.; Deinhardt, K.; West, J. Asymmetric Confinement for Defining Outgrowth Directionality. Lab. Chip 2019, 19 (8), 1484–1489. 10.1039/C9LC00078J.

(50) Girardin, S.; Clément, B.; J. Ihle, S.; Weaver, S.; B. Petr, J.; C. Mateus, J.; Duru, J.; Krubner, M.; Forró, C.; Ruff, T.; Fruh, I.; Müller, M.; Vörös, J. Topologically Controlled Circuits of Human iPSC-Derived Neurons for Electrophysiology Recordings. Lab. Chip 2022, 22 (7), 1386–1403. 10.1039/D1LC01110C.

(51) Smeal, R. M.; Rabbitt, R.; Biran, R.; Tresco, P. A. Substrate Curvature Influences the Direction of Nerve Outgrowth. Ann. Biomed. Eng. 2005, 33 (3), 376–382. 10.1007/s10439-005-1740-z.

(52) Francisco, H.; Yellen, B. B.; Halverson, D. S.; Friedman, G.; Gallo, G. Regulation of Axon Guidance and Extension by Three-Dimensional Constraints. Biomaterials 2007, 28 (23), 3398–3407. 10.1016/j.biomaterials.2007.04.015.

(53) Dickson, B. J. Molecular Mechanisms of Axon Guidance. Science 2002, 298 (5600), 1959–1964. 10.1126/science.1072165.

(54) Polleux, F.; Ince-Dunn, G.; Ghosh, A. Transcriptional Regulation of Vertebrate Axon Guidance and Synapse Formation. Nat. Rev. Neurosci. 2007, 8 (5), 331–340. 10.1038/nrn2118.

(55) Ballini, M.; Müller, J.; Livi, P.; Chen, Y.; Frey, U.; Stettler, A.; Shadmani, A.; Viswam, V.; Jones, I. L.; Jäckel, D.; Radivojevic, M.; Lewandowska, M. K.; Gong, W.; Fiscella, M.; Bakkum, D. J.; Heer, F.; Hierlemann, A. A 1024-Channel CMOS Microelectrode Array With 26,400 Electrodes for Recording and Stimulation of Electrogenic Cells In Vitro. IEEE J. Solid-State Circuits 2014, 49 (11), 2705–2719. 10.1109/JSSC.2014.2359219.

(56) Huh, D.; Matthews, B. D.; Mammoto, A.; Montoya-Zavala, M.; Hsin, H. Y.; Ingber, D. E. Reconstituting Organ-Level Lung Functions on a Chip. Science 2010, 328 (5986), 1662–1668. 10.1126/science.1188302.

(57) Maoz, B. M.; Herland, A.; FitzGerald, E. A.; Grevesse, T.; Vidoudez, C.; Pacheco, A. R.; Sheehy, S. P.; Park, T.-E.; Dauth, S.; Mannix, R.; Budnik, N.; Shores, K.; Cho, A.; Nawroth, J. C.; Segrè, D.; Budnik, B.; Ingber, D. E.; Parker, K. K. A Linked Organ-on-Chip Model of the Human Neurovascular Unit Reveals the Metabolic Coupling of Endothelial and Neuronal Cells. Nat. Biotechnol. 2018, 36 (9), 865–874. 10.1038/nbt.4226.

(58) Low, L. A.; Mummery, C.; Berridge, B. R.; Austin, C. P.; Tagle, D. A. Organs-on-Chips: Into the next Decade. Nat. Rev. Drug Discov. 2021, 20 (5), 345–361. 10.1038/s41573-020-0079-3.

(59) Macarthur, C. C.; Xue, H.; Van Hoof, D.; Lieu, P. T.; Dudas, M.; Fontes, A.; Swistowski, A.; Touboul, T.; Seerke, R.; Laurent, L. C.; Loring, J. F.; German, M. S.; Zeng, X.; Rao, M. S.; Lakshmipathy, U.; Chesnut, J. D.; Liu, Y. Chromatin Insulator Elements Block Transgene Silencing in Engineered Human Embryonic Stem Cell Lines at a Defined Chromosome 13 Locus. Stem Cells Dev. 2012, 21 (2), 191–205. 10.1089/scd.2011.0163.

(60) Vulić, K.; Amos, G.; Ruff, T.; Kasm, R.; Ihle, S. J.; Küchler, J.; Vörös, J.; Weaver, S. Impact of Microchannel Width on Axons for Brain-on-Chip Applications. Lab. Chip 2024. 10.1039/D4LC00440J.

(61) Sébastien, M.; Paquette, A. L.; Prowse, E. N. P.; Hendricks, A. G.; Brouhard, G. J. Doublecortin Restricts Neuronal Branching by Regulating Tubulin Polyglutamylation. Nat. Commun. 2025, 16 (1), 1749. 10.1038/s41467-025-56951-2.

(62) Yoon, S.-J.; Elahi, L. S.; Pașca, A. M.; Marton, R. M.; Gordon, A.; Revah, O.; Miura, Y.; Walczak, E. M.; Holdgate, G. M.; Fan, H. C.; Huguenard, J. R.; Geschwind, D. H.; Pașca, S. P. Reliability of Human Cortical Organoid Generation. Nat. Methods 2019, 16 (1), 75–78. 10.1038/s41592-018-0255-0.

(63) Duru, J.; Maurer, B.; Giles Doran, C.; Jelitto, R.; Küchler, J.; Ihle, S. J.; Ruff, T.; John, R.; Genocchi, B.; Vörös, J. Investigation of the Input-Output Relationship of Engineered Neural Networks Using High-Density Microelectrode Arrays. Biosens. Bioelectron. 2023, 239, 115591. 10.1016/j.bios.2023.115591.

(64) Radivojevic, M.; Franke, F.; Altermatt, M.; Müller, J.; Hierlemann, A.; Bakkum, D. J. Tracking Individual Action Potentials throughout Mammalian Axonal Arbors. eLife 2017, 6, e30198. 10.7554/eLife.30198.

(65) Molen, T. van der; Lim, M.; Bartram, J.; Cheng, Z.; Robbins, A.; Parks, D. F.; Petzold, L. R.; Hierlemann, A.; Haussler, D.; Hansma, P. K.; Tovar, K. R.; Kosik, K. S. RT-Sort: An Action Potential Propagation-Based Algorithm for Real Time Spike Detection and Sorting with Millisecond Latencies. PLOS ONE 2024, 19 (12), e0312438. 10.1371/journal.pone.0312438.

(66) Bakkum, D. J.; Frey, U.; Radivojevic, M.; Russell, T. L.; Müller, J.; Fiscella, M.; Takahashi, H.; Hierlemann, A. Tracking Axonal Action Potential Propagation on a High-Density Microelectrode Array across Hundreds of Sites. Nat. Commun. 2013, 4, 2181. 10.1038/ncomms3181.

(67) Nowakowski, T. J.; Bhaduri, A.; Pollen, A. A.; Alvarado, B.; Mostajo-Radji, M. A.; Di Lullo, E.; Haeussler, M.; Sandoval-Espinosa, C.; Liu, S. J.; Velmeshev, D.; Ounadjela, J. R.; Shuga, J.; Wang, X.; Lim, D. A.; West, J. A.; Leyrat, A. A.; Kent, W. J.; Kriegstein, A. R. Spatiotemporal Gene Expression Trajectories Reveal Developmental Hierarchies of the Human Cortex. Science 2017, 358 (6368), 1318–1323. 10.1126/science.aap8809.

(68) Avitan, L.; Stringer, C. Not so Spontaneous: Multi-Dimensional Representations of Behaviors and Context in Sensory Areas. Neuron 2022, 110 (19), 3064–3075. 10.1016/j.neuron.2022.06.019.

(69) Cai, H.; Ao, Z.; Tian, C.; Wu, Z.; Liu, H.; Tchieu, J.; Gu, M.; Mackie, K.; Guo, F. Brain Organoid Reservoir Computing for Artificial Intelligence. Nat. Electron. 2023, 6 (12), 1032–1039. 10.1038/s41928-023-01069-w.

(70) Robbins, A.; Schweiger, H. E.; Hernandez, S.; Spaeth, A.; Voitiuk, K.; Parks, D. F.; Molen, T. van der; Geng, J.; Cline, I.; Kosik, K. S.; Salama, S. R.; Sharf, T.; Mostajo-Radji, M. A.; Haussler, D.; Teodorescu, M. Goal-Directed Learning in Cortical Organoids. Cell Rep. 2026, 45 (2). 10.1016/j.celrep.2026.116984.

(71) Nguyen, C. V.; Martija, A.; Nano, P. R.; Soto, J. A.; Jaklic, D. C.; Mil, J.; White, R.; Martin, J. M.; Rana, D.; Geschwind, D. H.; Bhaduri, A. Thalamic NRXN1-Mediated Input to Human Cortical Progenitors Drives Upper Layer Neurogenesis. bioRxiv April 25, 2025, p 2025.04.25.650717. 10.1101/2025.04.25.650717.

(72) Patton, M. H.; Thomas, K. T.; Bayazitov, I. T.; Newman, K. D.; Kurtz, N. B.; Robinson, C. G.; Ramirez, C. A.; Trevisan, A. J.; Bikoff, J. B.; Peters, S. T.; Pruett-Miller, S. M.; Jiang, Y.; Schild, A. B.; Nityanandam, A.; Zakharenko, S. S. Synaptic Plasticity in Human Thalamocortical Assembloids. Cell Rep. 2024, 43 (8). 10.1016/j.celrep.2024.114503.

(73) Chilton, J. K. Molecular Mechanisms of Axon Guidance. Dev. Biol. 2006, 292 (1), 13–24. 10.1016/j.ydbio.2005.12.048.

(74) Birtele, M.; Lancaster, M.; Quadrato, G. Modelling Human Brain Development and Disease with Organoids. Nat. Rev. Mol. Cell Biol. 2025, 26 (5), 389–412. 10.1038/s41580-024-00804-1.

(75) Sun, Y.; Ikeuchi, Y.; Guo, F.; Hyun, I.; Ming, G.; Fu, J. Bioengineering Innovations for Neural Organoids with Enhanced Fidelity and Function. Cell Stem Cell 2025, 32 (5), 689–709. 10.1016/j.stem.2025.03.014.

(76) Sakaguchi, H.; Ozaki, Y.; Ashida, T.; Matsubara, T.; Oishi, N.; Kihara, S.; Takahashi, J. Self-Organized Synchronous Calcium Transients in a Cultured Human Neural Network Derived from Cerebral Organoids. Stem Cell Rep. 2019, 13 (3), 458–473. 10.1016/j.stemcr.2019.05.029.

(77) Aggarwal, A.; Liu, R.; Chen, Y.; Ralowicz, A. J.; Bergerson, S. J.; Tomaska, F.; Mohar, B.; Hanson, T. L.; Hasseman, J. P.; Reep, D.; Tsegaye, G.; Yao, P.; Ji, X.; Kloos, M.; Walpita, D.; Patel, R.; Mohr, M. A.; Tillberg, P. W.; Looger, L. L.; Marvin, J. S.; Hoppa, M. B.; Konnerth, A.; Kleinfeld, D.; Schreiter, E. R.; Podgorski, K. Glutamate Indicators with Improved Activation Kinetics and Localization for Imaging Synaptic Transmission. Nat. Methods 2023, 20 (6), 925–934. 10.1038/s41592-023-01863-6.

(78) Marvin, J. S.; Shimoda, Y.; Magloire, V.; Leite, M.; Kawashima, T.; Jensen, T. P.; Kolb, I.; Knott, E. L.; Novak, O.; Podgorski, K.; Leidenheimer, N. J.; Rusakov, D. A.; Ahrens, M. B.; Kullmann, D. M.; Looger, L. L. A Genetically Encoded Fluorescent Sensor for in Vivo Imaging of GABA. Nat. Methods 2019, 16 (8), 763–770. 10.1038/s41592-019-0471-2.

(79) Birtele, M.; Del Dosso, A.; Xu, T.; Nguyen, T.; Wilkinson, B.; Hosseini, N.; Nguyen, S.; Urenda, J.-P.; Knight, G.; Rojas, C.; Flores, I.; Atamian, A.; Moore, R.; Sharma, R.; Pirrotte, P.; Ashton, R. S.; Huang, E. J.; Rumbaugh, G.; Coba, M. P.; Quadrato, G. Non-Synaptic Function of the Autism Spectrum Disorder-Associated Gene SYNGAP1 in Cortical Neurogenesis. Nat. Neurosci. 2023, 26 (12), 2090–2103. 10.1038/s41593-023-01477-3.

