## Supplementary Figures 1-6 for "Spatially defined axonal guidance in neural organoids with micropatterned microfluidic channels"

Supplementary Information for

### Engineering in vitro neurite outgrowth with directional microstructures

Supplemental Figure 1: Microstructure Fabrication

Supplemental Figure 2: Design Iterations

Supplemental Figure 3: Control Microstructures

Supplemental Figure 4: Electrophysiology impedance scanning source/target selection

Supplemental Figure 5: Comparison of spike frequency distributions in asymmetric versus straight channel groups

Supplemental Figure 6: Firing rate distribution along microchannel distance at 5% bins

**This PDF file includes:**

Supplementary Figures 1 to # 6

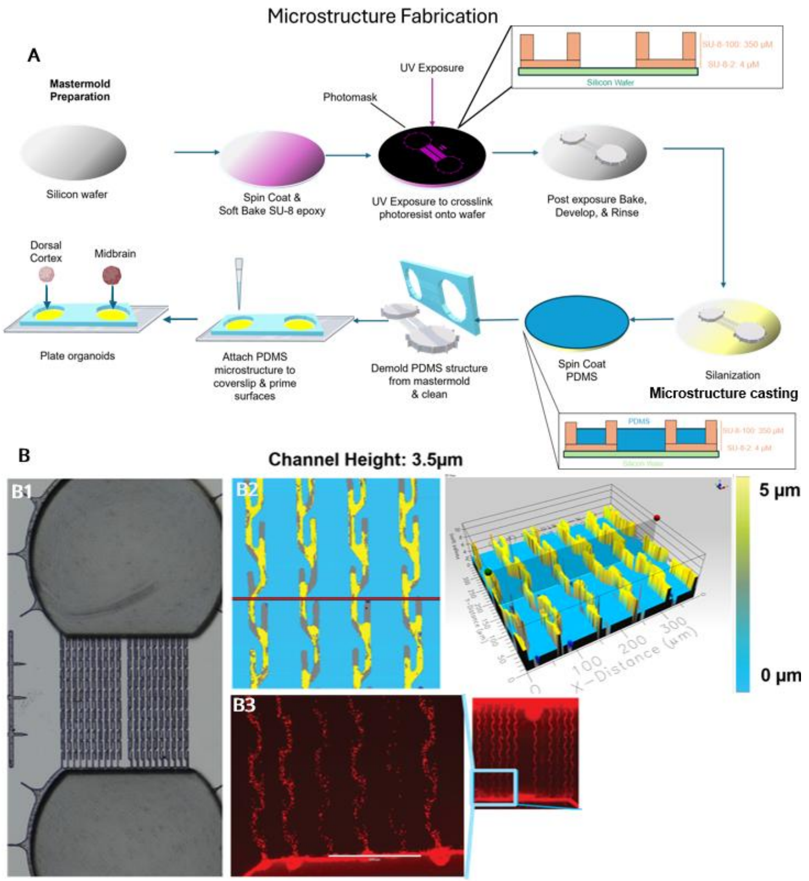


**Supplementary Figure 1: Fabrication of microchannels (Supp.1A)** Fabrication workflow of photo patterned mastermold **(Supp. B1-B3)** Directoid mastermold and feature testing. **B1:** Brightfield image of fabricated mastermold. **B2**: Confirmed channel height at 3.5 µm by 2D profilometer. **B3**: After PDMS microstructure casting, removal of uncured PDMS with overnight ethyl acetate wash and priming of channels on coverslip, 10nm latex beads are passed a through sacrificial microstructure channels under vacuum to ensure open channels.

**Commented [1]:** Nice figure! Scale bars are missing and/or hard to read

Supplemental Figure 2: Design iterations

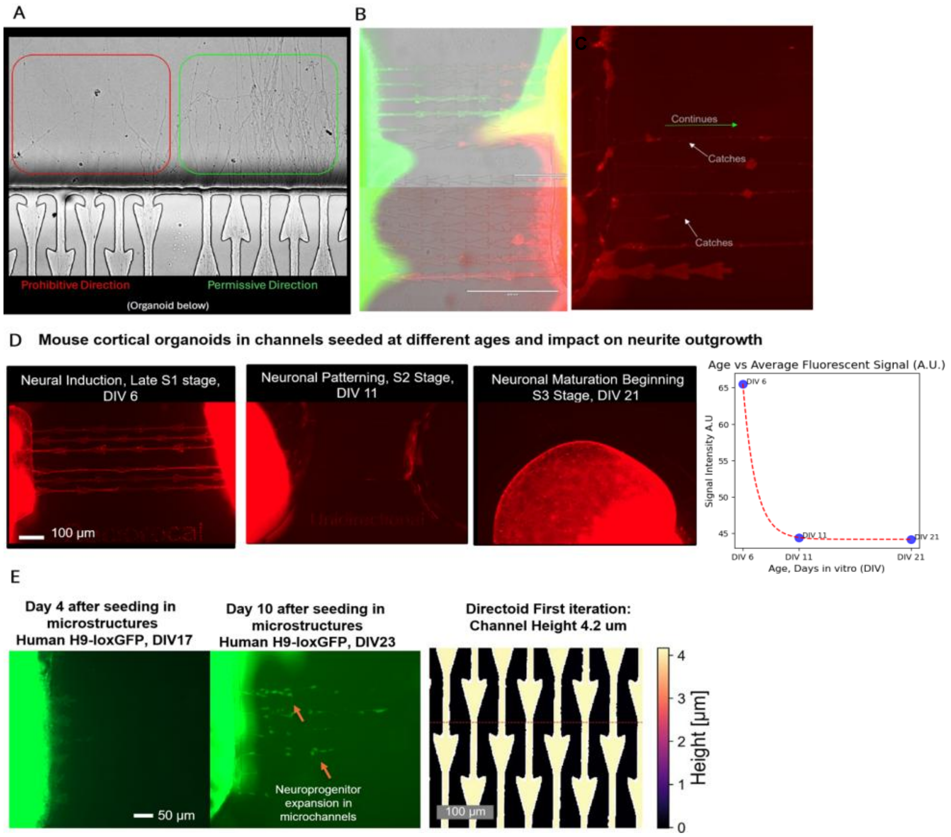


**Supplementary Figure 2. Design Iterations Reveal Structural and Developmental Constraints on Organoid Integration into Microchannels.** (**A**) Brightfield image of early asymmetric microchannel design, adapted from 2D axon guidance literature, intended to bias axon outgrowth. Axons showed preferential growth in the permissive direction (green), but prohibitive regions (red) still allowed occasional entry, indicating insufficient directionality enforcement. (**B**) Overlay of GFP+ and mCherry+ axons in co-cultured organoids demonstrates lack of directional segregation, with widespread invasion across both channel orientations.(**C**) High-resolution fluorescence image of axon behavior within channels shows growth cones either continuing through permissive zones (green arrow) or catching at junctions (white arrows), highlighting structural inefficiencies in early designs. Height of channels is 5.1µm(**D**) Mouse cortical organoids seeded at 6, 11, and 21 days in vitro (DIV) into various microchannel formats (reciprocal, unidirectional, open well) showed robust axon entry only at early stages. At DIV21, all organoids were infected with pAAV-hSyn-mScarlet (AAV1) and images taken at

**Commented [2]:** Please specify the channel heights here. This was a earlier iteration with taller channel heights, and we see cells migrating. We want to make the point the ~4 um channel heights prevent this (Fig 1 and Suppl fig 1)

**Commented [AC3R2]:** Do Days post seeding and DIV, not S-media (Seeded at XDIV N, DIV Y when image taken)

approximately DIV30. We observed channel engagement sharply declining with organoid age, as illustrated in the adjacent schematic, emphasizing the narrow developmental window for effective axon guidance in murine models. (**E**) Human organoids demonstrated broader temporal plasticity. Human organoids were also initially seeded soon after the embryoid body stage, at DIV15, to mirror similar timelines observed in mouse prototypes. However, at DIV17 (Day 4 post-seeding), while neurite outgrowth was too early to be observed, tissue migration was observed in channels. By DIV23, this tissue was confirmed to be neuroprogenitor migration into microchannels (arrows), followed by in situ differentiation. This emphasized the need to decouple organoid age from seeding age for optimal integration and structural patterning. Furthermore, neuroprogenitor migration into microchannels also necessitated decreasing channel height in order minimize neuroprogenitor migration, especially for organoid types where neuroprogenitor pools persist for longer periods of time, such as human cortical organoids.

**Commented [4]:** We need time points for how old the organoids are and when they we seeded in these images. Can you generate a real graph of this bevavior?

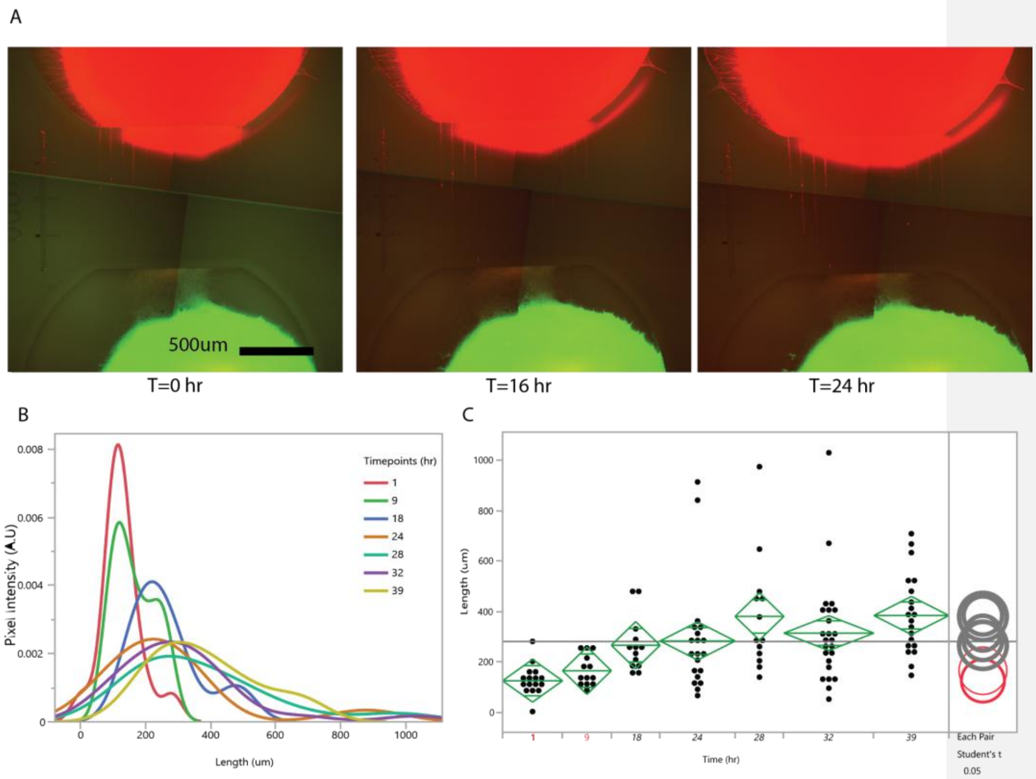


**Supplementary Figure 3. Live Imaging of axon migration in control microchannels**

A. Time-course images of axon migration of H9-cag-mCherry dorsal forebrain axon migration at 0, 16, and 24 hours.

**Commented [5]:** Scale bars :)
Also, make sure that all SI figures are referenced in the text [results section and/or materials]. I.e. the observation that the younger the organoid, the more axonal outgrowth is quite a nice finding and should definitely be mentioned in the main text and referencing the SI

B. Normalized pixel intensity distribution profiles along the migration path at multiple time points (0-39 hours), showing the progressive extension and advancement of the fluorescent signals over time.
C. Quantitative analysis of axon migration length versus time, with individual data points and box plots showing the distribution of migration distances at each time point. Student's t-test. P-value < 0.05 demonstrates significant changes in axon positioning over the experimental duration, illustrating the temporal dynamics of axon growth in control microstructures.

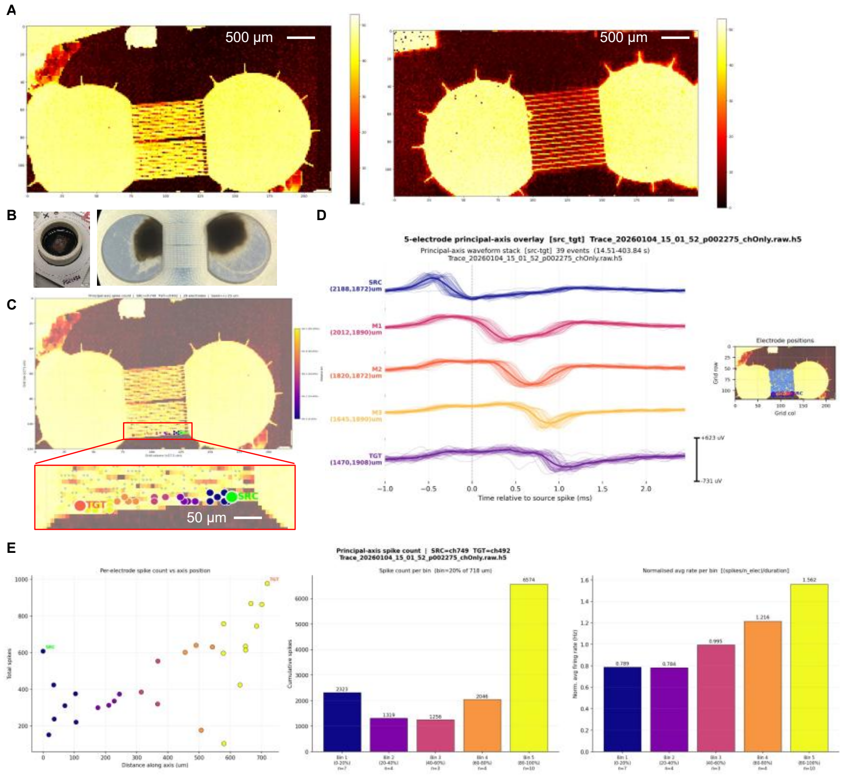


**Supplementary Figure 4. Electrophysiology impedance scanning source/target selection**

A. Impedance Scan: After microstructure attachment and surface laminin treatment, a threshold voltage is applied to the HD-MEA to generate differential impedances, between sealed and unsealed portions of the microchip and identify electrodes uncovered by PDMS. Selected electrodes during passive recordings were routed to available amplifiers and were used to create recording configurations over the microchannels. (Left) microstructure is an asymmetric directional microstructure (directoid) (Right) microstructure is an straight connectoid microstructure.

B. Organoid Attachment: After impedance scanning, organoids are attached to chips. A sacrificial microstructure is placed on glass coverslips to confirm outgrowth of organoid batch into microchannels visually before stochastic action potentials appear.

C. After recording is performed, our recording toolkit enables initial peak finding and manual annotation of microstructure channels can be performed by selecting a Source electrode (Green) and a Target electrode (Orange). Principal axis band with a 1 pixel search radius captures intermediate electrodes, bins, and labels them by quintile.

D. **SpikePathTool, Branch B**: After Source/Target and principal axis band has been selected script allows either manual or automatic selection of intermediate electrodes and display position and spikes that occur during 1-2ms firing window between Source and Target electrode respectively. Latency plots pull and plot >30 Source/Target windows from raw data traces and overlays 2.5 ms windows from all selected electrodes.

E. **SpikePathTool, Branch A**: SpikePathTool will also output overall total spikes for all electrodes selected as a scatter plot of Total Spikes vs Distance along the principal axis (Left), Bar plot of the Total Number of Spikes separated by spatial bin (Middle), and Bar plot of Bin grouped normalized average firing rate.

Supplement 5 Comparison of spike frequency at 25% & 75% quartile of channel by microstructure, Directoids

**A**

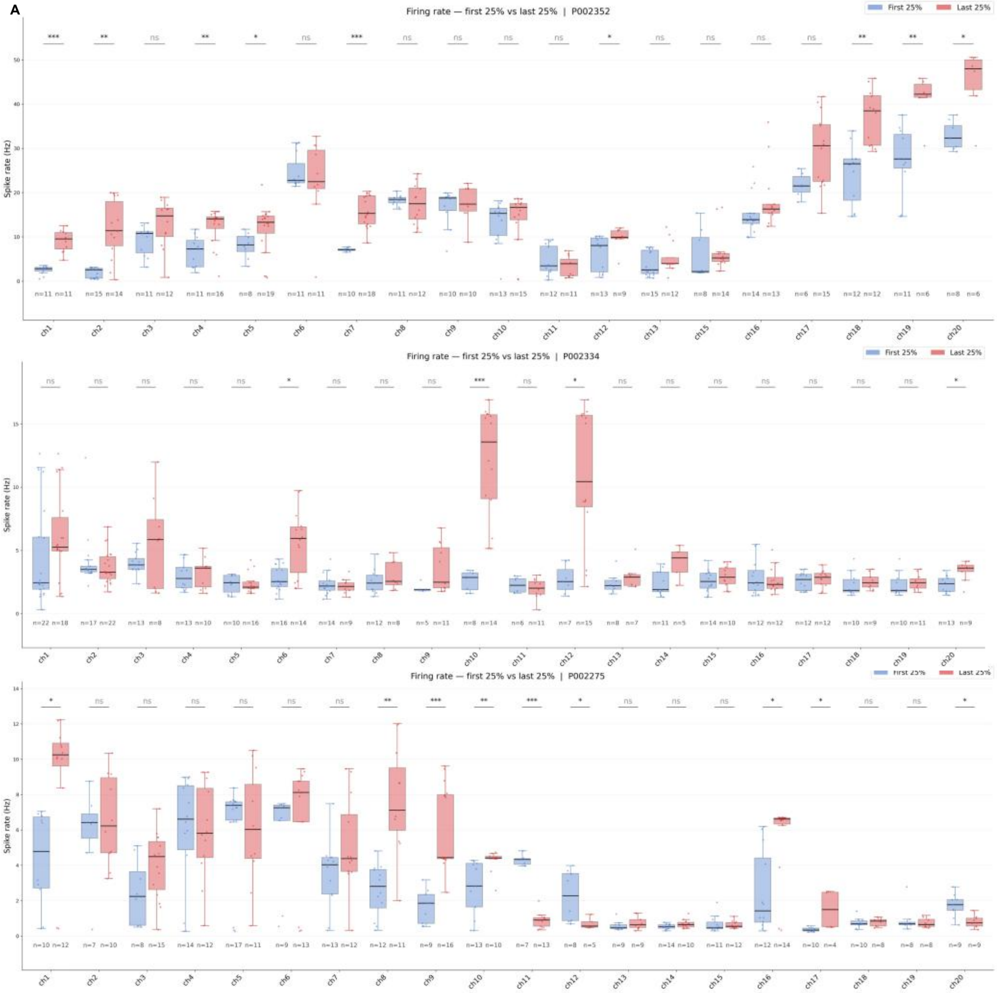

Supplement 5 Comparison of spike frequency at 25% & 75% quartile of channel by microstructure, Connectoids

**B**

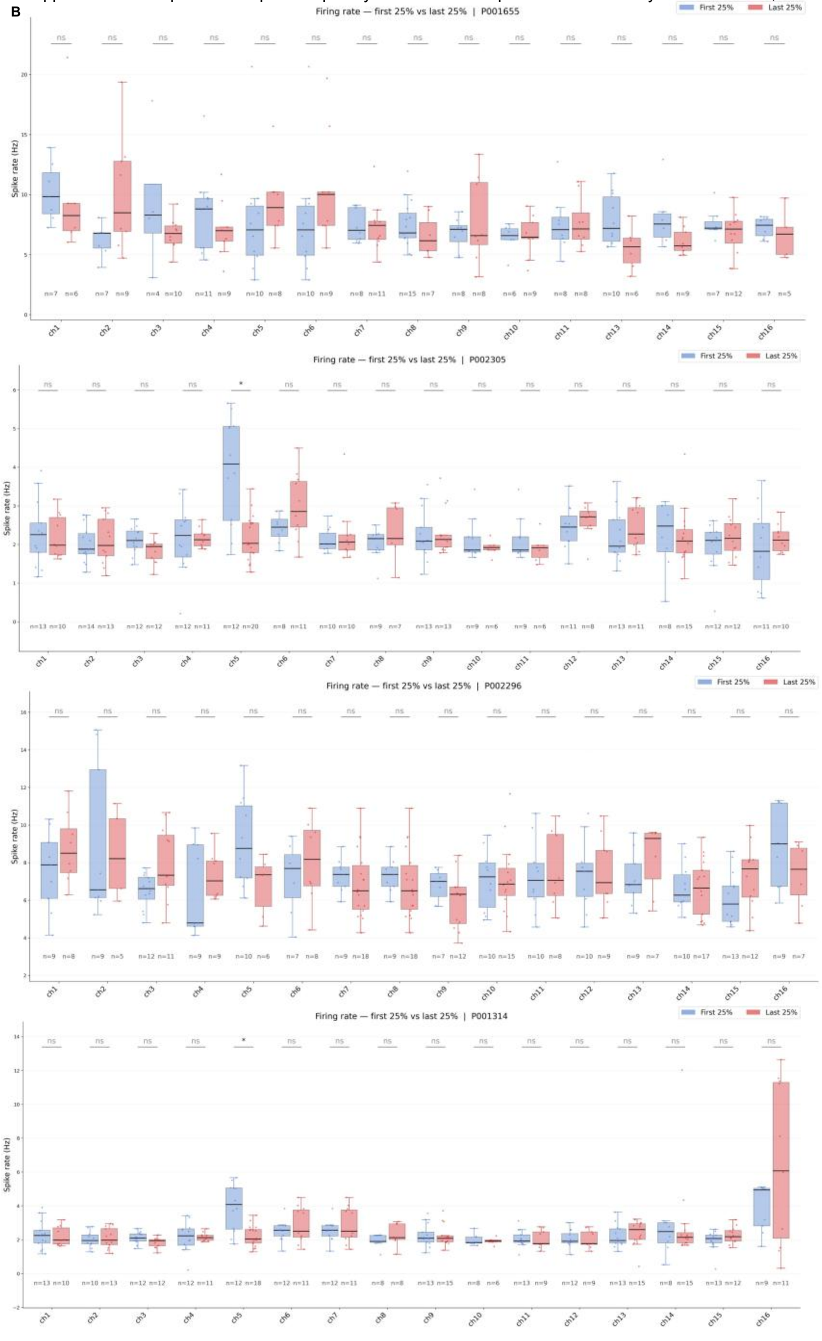

**Supplementary Figure 5. Comparison of spike frequency distributions in asymmetric versus straight channel groups.** Box-and-whisker plots showing spike frequency (Hz) across channels, comparing activity within the first 25% and last 25% of microchannel distance along a selected principal axis. Asymmetric channels in **A** demonstrate multiple channels with consistent, asymmetric spike propagation along multiple microchannel paths, while undirected channels in **B** shows straight channels, which lack a dominant propagation direction and exhibit more symmetric, diffuse activity. Blue and red indicate early and late microchannel segments, respectively. Asymmetric channels exhibit more coherent temporal modulation of spike frequency, whereas undirected channels show weaker and more variable changes. Paired regions of microchannels that demonstrate significant differences in firing rate (ANOVA, $p<0.05$) were denoted as the fraction of channels that demonstrated significant changes in firing rate in Figure5E

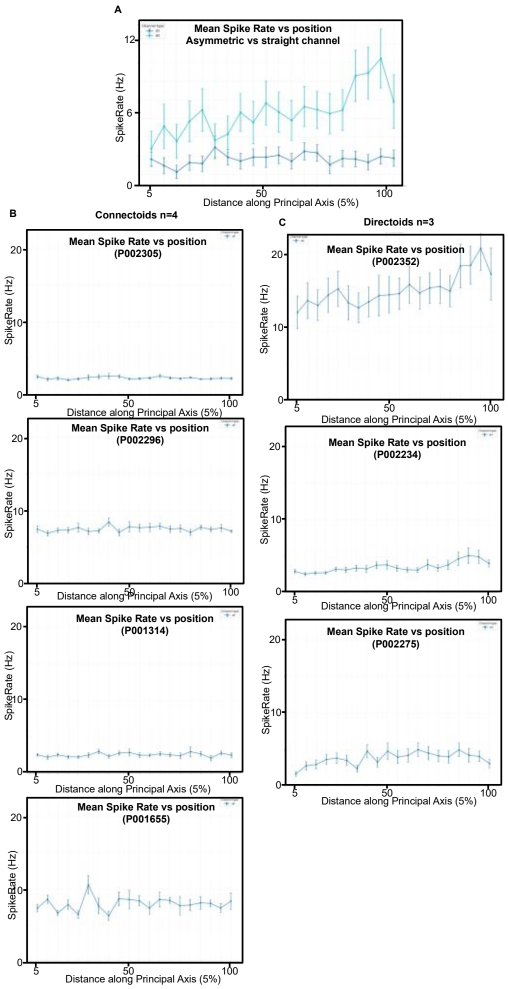


**Supplemental Figure 6: Firing rate distribution along microchannel distance at 5% bins:**

A) Comparison of mean firing rates between both asymmetric directoid microstructures and straight connectoid microstructures binned by 5% intervals along microchannel length.
B) Line plot of mean firing rate and SEM of all microchannels for each straight channel connectoid microstructure binned by 5%distances along principal axis for each straight microchannel connectoid microstructure
C) Line plot of mean firing rate and SEM of all microchannels asymmetric channel directoid microstructure binned by 5%distances along principal axis for each asymmetric microchannel directoid microstructure
